# MK2 deficiency decreases mortality during the inflammatory phase after myocardial infarction in mice

**DOI:** 10.1101/2023.03.06.531384

**Authors:** Joëlle Trépanier, Sherin A. Nawaito, Pramod Sahadevan, Fatiha Sahmi, Natacha Duquette, Danielle Gélinas, Marc-Antoine Gillis, Yanfen Shi, Cynthia Torok, Marie-Élaine Clavet-Lanthier, Gaestel Matthias, Martin G. Sirois, Jean-Claude Tardif, Bruce G. Allen

## Abstract

**Background:** Altering the onset, intensity, or duration of inflammation can impact the recovering heart’s structure and function following myocardial infarction (MI). Substrates of MAP kinase-activated protein kinase 2 (MK2) include proteins that regulate the stability of AU-rich transcripts, including those of several pro-inflammatory cytokines. This study was to determine if MK2-deficiency impaired the inflammatory phase of post-MI wound repair.

**Methods and Results:** Myocardial infarctions were induced by permanent ligation of the left anterior descending coronary artery in 12-week-old male MK2^+/+^ and MK2^-/-^ littermate mice. Five days post-MI, survival was 100% in MI-MK2^-/-^ (n = 20) and 79% in MI-MK2^+/+^ mice (n = 29; Mandel-Cox test: *P* < 0.05). Area at risk and infarct size were similar. Echocardiographic imaging revealed that both systolic and diastolic LV diameters were greater in MI-MK2^+/+^ than MI-MK2^-/-^ mice. MK2-deficiency did not affect the increase in wall motion score index. Infiltration of neutrophils or monocytes did not differ significantly. Cytokine and chemokine transcripts were quantified in infarcted and non-infarcted LV tissue using qPCR arrays (QIAGEN). Three days post-MI, *Ifna2* was increased and *Il16* was decreased in infarcted tissue from MK2^-/-^ hearts, compared with infarcted MK2^+/+^ tissue, whereas in the non-infarcted MK2^-/-^ myocardium *Il27* increased and *Tnfsf11*, *Ccl3*, and *Il1rn* were decreased. Five days post-MI, *Ctf16* and *Il10* increased in infarcted MK2^-/-^ tissue whereas in the non-infarcted MK2^-/-^ myocardium *Ccl9, Nodal, and Xcl2* increased and *Il15* decreased.

**Conclusions:** The findings of this study suggest MK2-deficiency is an advantage during the inflammatory phase of cardiac wound repair post-MI.

**Clinical Perspective:** **What is new?**

-The effects of MAP kinase-activated protein kinase 2 (MK2) deficiency on survival, cardiac structure and function, and the inflammatory phase of wound healing following myocardial infarction were assessed using a constitutive, pan MK2-null mouse model.

-MK2-deficiency reduced mortality but did not alter area at risk or infarct size post-myocardial infarction. Inflammatory cell infiltration was also unaffected.

-MK2-deficiency altered the abundance of several cytokines (increased, decreased) in infarcted and non-infarcted myocardium post-MI.

**What are the clinical implications?**

-The initial phase of wound repair post-MI involves inflammation.

-The risk of damage to the myocardium and mortality may be reduced by inhibition of MK2 activity during the inflammatory phase of wound healing post-MI.

## Introduction

Myocardial infarction (MI) results from the partial or total occlusion of a coronary artery, which leads to necrosis and irreversible damage to the heart ^1–4^. Specifically, the lack of oxygen and nutrients causes a loss of cardiomyocytes (CM). As myocytes cannot replicate at the scale required to maintain muscle mass, they must be replaced by formation of a scar ^1, 2^. Inflammation is the first phase of the post-MI healing process and is key to the subsequent healing phases, proliferation and maturation ^3, 5^, as it orchestrates immune cell recruitment and fibroblast activation to myofibroblasts as well as the clearance of extracellular matrix (ECM) and cell debris, all of which are crucial to the formation of new scar tissue ^1, 3, 5–7^. Initially, necrosis results in the release of various signaling molecules such as TNF-α, IL-1 and other damage-associated molecular patterns (DAMPs), which trigger the activation of the immune system ^6, 8, 9^. Neutrophils are the first immune cells attracted in the ischemic area ^10^. They then actively participate in the recruitment of circulating monocytes and their activation to pro-inflammatory M1 macrophages ^10^. M1 macrophages and neutrophils clear the cellular and ECM debris by both phagocytosis and secretion of various proteases ^3, 4, 6^. They also secrete numerous cytokines including osteopontin (SPP1), granulocyte-macrophage colony-stimulating factor (CSF2), and adhesion molecules that participate in amplifying the recruitment of additional activated cells ^3, 6, 11–17^. This cascade of signals promotes fibroblast migration into the infarct, where they activate to myofibroblasts and secrete ECM components ^3, 4, 6^. Through this process of reparative fibrosis, a collagen scar is formed that both replaces the lost myocytes and permits the affected myocardium to withstand the pressure of the contracting heart without rupturing ^3, 5, 10, 18^.

Since inflammation initiates the post-MI healing process, alteration of the onset, intensity or duration of this response can have dramatic consequences on the heart’s structure and function following MI. A robust inflammatory response can induce excessive secretion of cytokines and proteases ^3^, resulting in additional myocyte loss and degradation of the ECM ^19–22^, further weakening the ventricular wall, leading to thinning of the wall and dilatation 3^, 19, 21–23^. In such cases, ventricular rupture is more likely as the ventricular wall is no longer able to withstand the pressure developed during cardiac contractions ^3, 19, 21–23^. Myocardial rupture could also occur if the inflammatory response is too weak to trigger reparative fibrosis ^24, 25^. In this case, the migration of fibroblasts into the ischemic area and activation to myofibroblasts would be insufficient, leaving the wall in a weakened state and prone to rupture ^24^. A prolonged inflammatory response can promote interstitial fibrosis formation, which would dramatically alter cardiac structure and function 3^, 5, 11, 23^, increasing LV wall stiffness and impairing myocyte synchronization ^24, 26–28^, leading to life-threatening conditions such as arrhythmias and heart failure ^3, 26, 28, 29^. Various mechanisms exist to resolve the inflammatory response in a timely manner, such as a shift from recruitment of Ly-6C^hi^ monocytes to Ly-6C^lo^ monocytes and macrophage M2 polarization ^30–32^. M2 macrophages secrete anti-inflammatory, pro-angiogenic and profibrotic factors such as IL-10 and TGF-β ^5, 6, 31, 33^. However, if these resolving mechanisms are hindered, inflammation will be prolonged ^27, 31^ and excessive neutrophil and M1 macrophage activity will lead to a scar expansion ^27, 31^. Thus, alterations in the coordinated events comprising the inflammatory phase of post-MI wound repair can have detrimental effects on the healing process that alter the structural integrity and functionality of the heart.

MAP kinase-activated protein kinase 2 (MK2) is a protein serine/threonine kinase that is activated by p38α and p38β MAPKs ^34–37^. The three-tiered classical MAPK cascade amplifies the signal initiated by extracellular stimuli, such as tissue damage or an infection, leading to an appropriate cellular response ^38–42^. These responses may include migration, proliferation, and differentiation, as well as regulation of the innate and adaptive immune system ^38–40, 43^. Many roles have been attributed to MK2, including remodeling of the actin cytoskeleton, apoptosis, regulation of transcription factors, and genomic stability ^36, 43–45^. MK2 is also a significant regulator of inflammation, as its inactivation is associated with decreased production of cytokines such as TNF-α, IL-1β, IL-6, IL-10, and INF-γ following a lipopolysaccharide (LPS) challenge in experimental models such as MK2-deficient mice, isolated splenocytes, and macrophages ^40, 43, 46–48^. MK2 regulates the stability of numerous pro-inflammatory cytokine transcripts through phosphorylation of RNA-binding proteins such as tristetraprolin (TTP), AU-binding factor 1 (AUF1), and human antigen R (HuR) ^43, 49^. Phosphorylation alters the affinity of these RNA binding proteins for the AU-rich elements in the 3’-UTRs of various mRNAs ^43, 49^. Protein binding to the 3’-UTR is a determinant of stability and thus alters the half-life of the mRNA ^43, 49^. In resting cells, TTP binding destabilizes target transcripts such as CSF2, IL-1, TNF-α, and IL-6, which continuously inhibits activation of the inflammatory response ^36, 43, 48, 50^. MK2-mediated phosphorylation of TPP at Ser-82 and Ser-178 reduces its affinity for its target mRNAs and allows HuR to bind and stabilize these transcripts ^43, 44, 48–51^. The resulting increase in transcript stability permits translation of various pro-inflammatory cytokines and ultimately triggers inflammation ^43, 44, 48–51^. Thus, MK2 deficiency in mice alters the inflammatory response and reduces mortality post-LPS challenge ^40, 43, 46–48, 52, 53^. *In vivo*, MK2-deficiency increases resistance to collagen-induced arthritis ^54^ and reduces atherosclerosis in hypercholesterolemic (*Ldlr*^-/-^:*Mk2*^-/-^) mice ^55^.

The acute, cardiomyocyte-specific activation of p38*α* in adult mice results in cardiomyocyte hypertrophy, interstitial fibrosis, contractile dysfunction, and mortality within one week ^53^. Although such studies suggest p38 would be a suitable target for drug development, several p38α inhibitors have failed as treatments against inflammatory diseases due to their high toxicity and lack of long-term efficacy ^43, 50, 56–58^. Furthermore, although targeted inactivation of p38α in platelets protected cardiac function after permanent ligation of the coronary artery ^59^, inactivation of p38α in cardiac fibroblasts led to 100% mortality in mice ^26^. Thus, downstream targets of p38, such as MK2, may serve as more suitable targets for drug development that would circumvent the issues associated with direct inhibition of one or more p38 isoform ^43, 56^. MK2-deficiency attenuates the hypertrophic effect and prevented the early mortality induced by cardiomyocyte-specific activation of p38 ^53^ whereas hypertrophy secondary to a chronic increase in afterload is delayed ^60^. In addition, MK2-deficiency prevents or delays the development of a diabetic cardiomyopathy in a mouse model of type 2 diabetes ^56, 61^. Since a proper inflammatory response is key to effective wound repair post-MI and MK2 is an important modulator in inflammation, this study was undertaken to assess the consequences of MK2 deficiency on mortality during the inflammatory phase of reparative fibrosis post-MI in mice.

## Material and Methods

Reagents for sodium dodecyl sulfate–polyacrylamide gel electrophoresis (SDS-PAGE), nitrocellulose membranes, and Bradford protein assay were from Bio-Rad Laboratories. Leupeptin and phenylmethylsulfonyl fluoride (PMSF) were from Roche Molecular Biochemicals. Rabbit polyclonal antibodies against p38α (# C-20) and MK2 (# 3042S) were from Santa Cruz Biotechnology and Cell Signaling Technology, respectively. Mouse monoclonal antibodies against GAPDH (glyceraldehyde 3-phosphate dehydrogenase) (# 4300) were from Ambion. Secondary antibodies conjugated with horseradish peroxidase were from Jackson ImmunoResearch Laboratories. Other reagents were either of analytic grade or the best grade available. The primers used for quantitative polymerase chain reaction (qPCR) reactions (**Table 1**) were produced by Invitrogen and their efficacy was previously demonstrated ^62^.

**Table 1.**
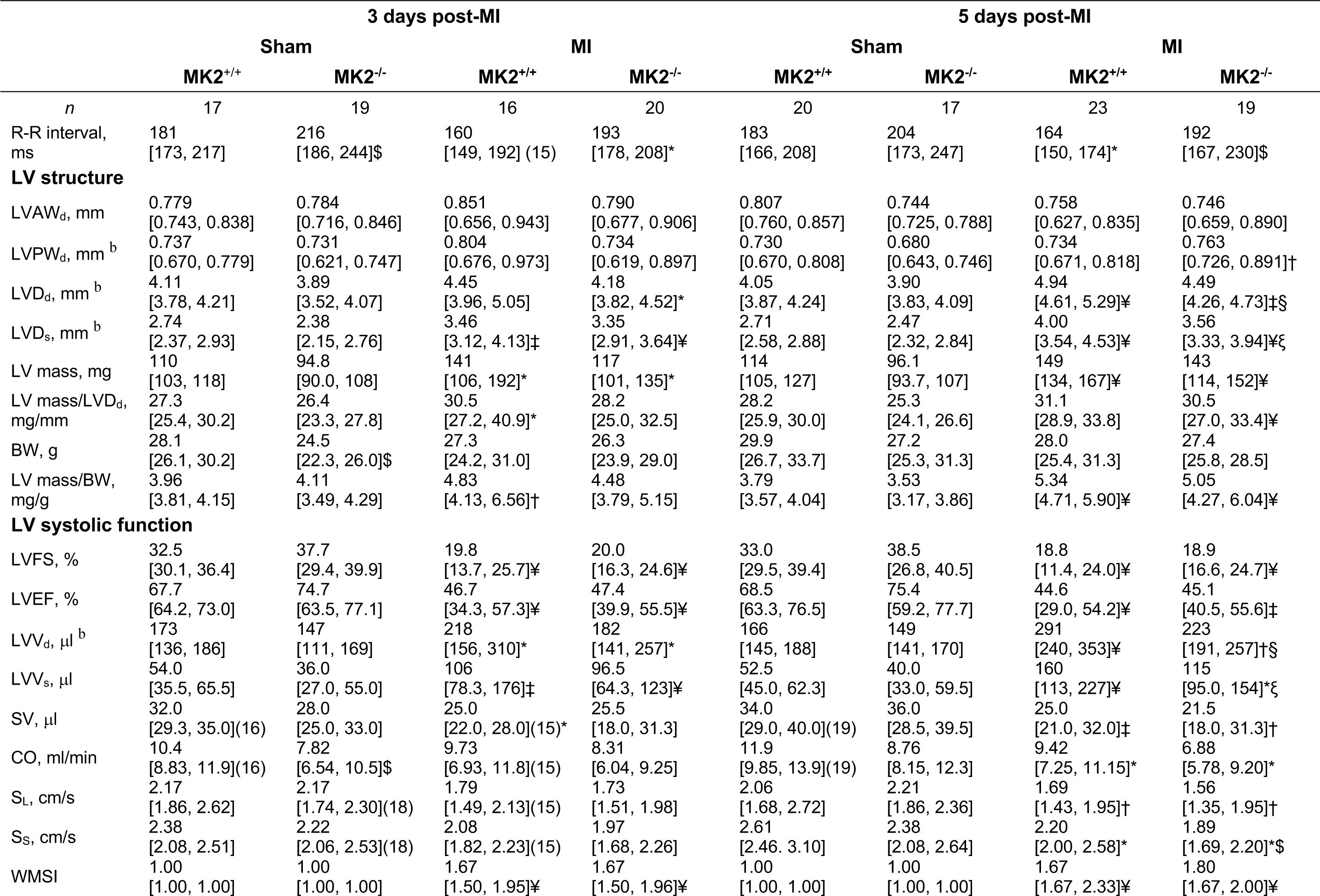
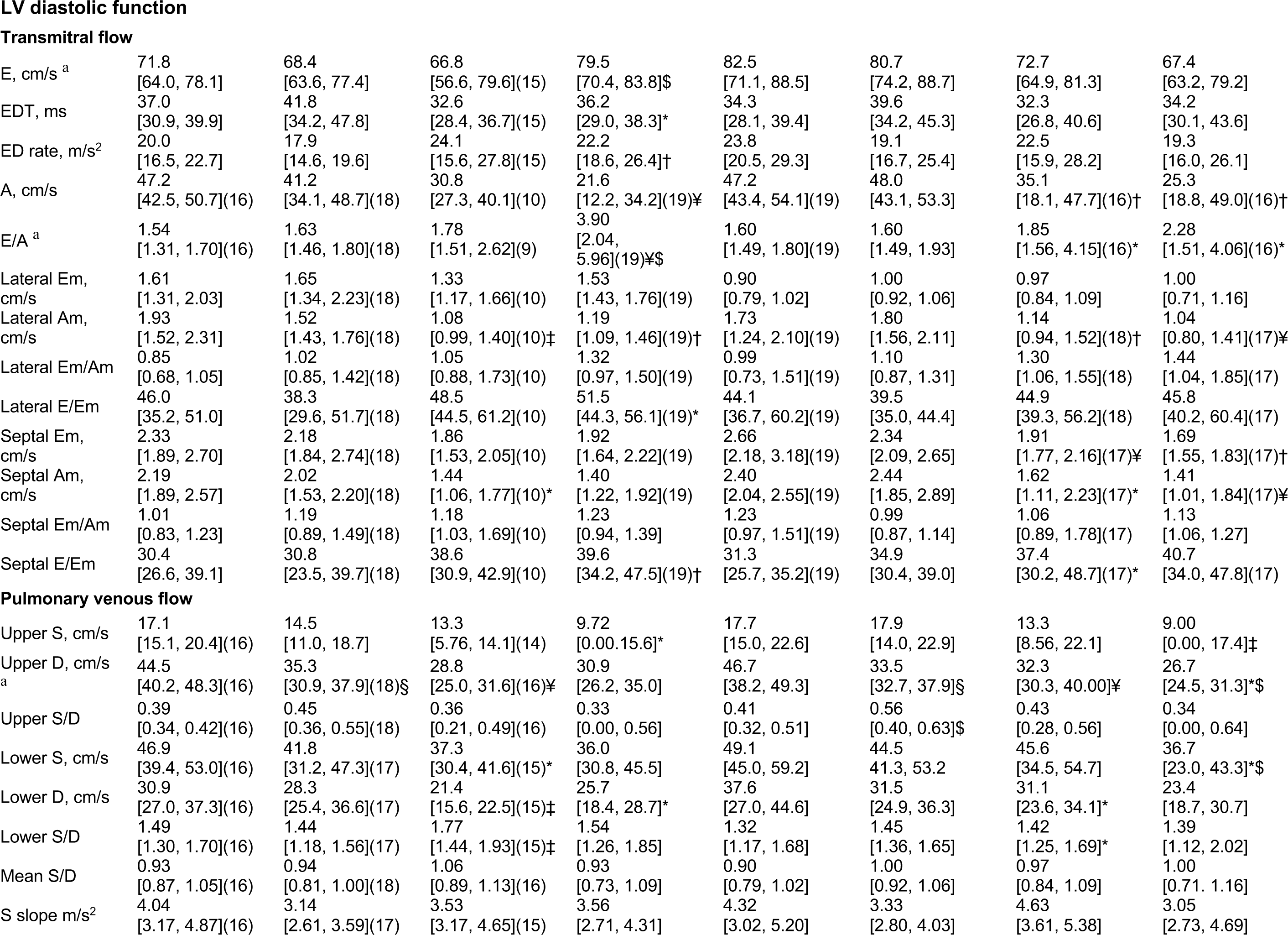
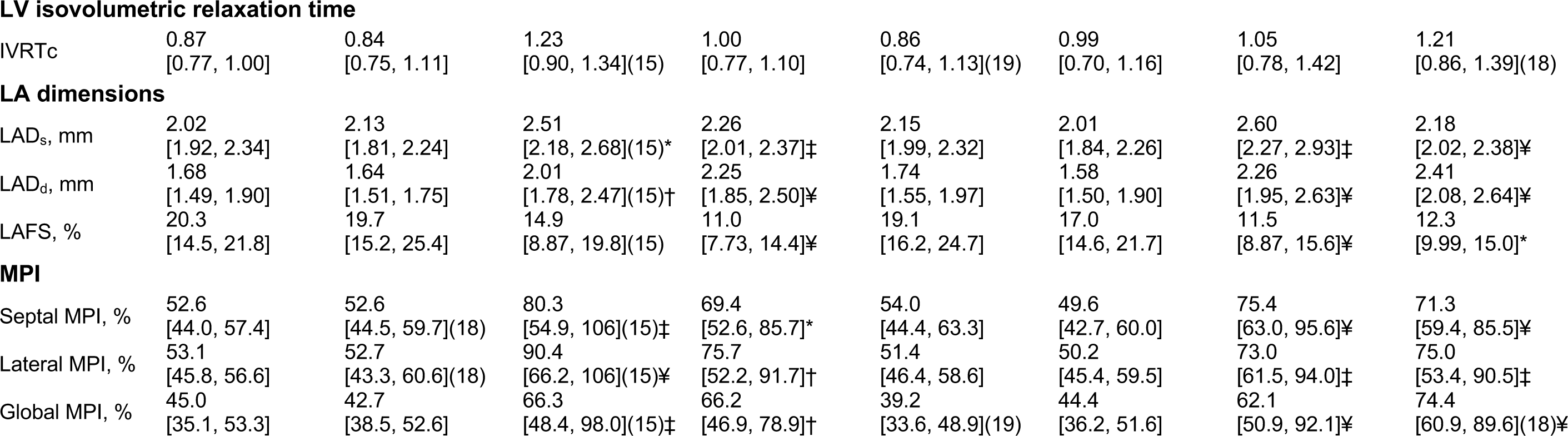
Echocardiography parameters of LV structure and function in 12-week-old MK2^+/+^ and MK2^-/-^ mice at 3 or 5 days post-MI. Data are reported as median [1^st^ quartile, 3^rd^ quartile]. The *n* numbers are as shown in the first row of the table unless otherwise indicated in parentheses. A indicates trans mitral flow late (atrial) filling velocity; Am, mitral annulus peak velocity during atrial diastolic filing; BW, body weight; CO, cardiac output; D, peak velocity during pulmonary venous diastolic flow; E, transmitral flow early filling velocity; ED, E wave deceleration; EDT, E wave deceleration time; Em, mitral annulus peak velocity during early diastolic filling; IVRT, isovolumic relaxation time; IVRTc: heart rate-corrected IVRT; LADd, left atrium dimension at end cardiac diastole; LADs, left atrium dimension at end cardiac systole; LAFS, left atrium fractional shortening; LV, left ventricular; LVAWd, left ventricular anterior wall thickness at end cardiac diastole; LVDd, left ventricular dimension at end cardiac diastole; LVDs, left ventricular dimension at end cardiac systole; LVEF, left ventricular ejection fraction; LVFS, left ventricular fractional shortening; LVPWd, left ventricular posterior wall thickness at end cardiac diastole; LVVd, left ventricular volume at end cardiac diastole; LVVs, left ventricular volume at end cardiac systole; MK2, mitogen-activated protein kinase–activated protein kinase-2; MPI, myocardial performance index; S, peak velocity during pulmonary venous systolic flow; SD, slope: pulmonary venous systolic flow decelerating slope; SL, basal lateral systolic velocity; SS, basal septal systolic velocity; SV, stroke volume; and WMSI, wall motion score index. Shapiro-Wilk tests for normality were performed for all data: if a lognormal distribution was indicated, data was log-transformed prior to statistical comparison by two-way ANOVA, including a factor for surgery (sham, MI), a factor for genotype (MK2^+/+^, MK2^-/-^), and a surgery x genotype interaction term. The ANOVA was followed by Tukey’s post hoc tests for multiple comparisons to compare means. *: *P* < 0.05 vs sham. †: *P* < 0.01 vs sham. ‡: *P* < 0.001 vs sham. ¥: *P* < 0.0001 vs sham. $: *P* < 0.05 vs MK2^+/+^. ξ: *P* < 0.01 vs MK2^+/+^. §: *P* < 0.001 vs MK2^+/+^. ^a^: *P* < 0.05 for surgery x genotype interaction 3-days post-MI. ^b^: *P* < 0.05 for surgery x genotype interaction 5-days post-MI.

### Mice

MK2-deficient mice (MK2^-/-^) were generated by insertion of a neomycin resistance gene, which contains an in-frame translation stop codon, into the exon containing the catalytic subdomains V and VI of the protein ^46^. The result is an inactive MK2 protein truncated at the active site. Twelve-week-old male MK2^+/+^ and MK2^-/-^ littermate mice were used for these experiments. MK2^-/-^ mice do not display any adverse physiological or behavioral defects ^36^. They are viable and fertile ^36^. All experiments were approved by the local ethics committee and performed according to the guidelines of the Canadian Council on Animal Care. The mice were housed in a specific pathogen-free facility maintained at a constant room temperature and with a 12-hour light/dark cycle.

### Myocardial infarction

Myocardial infarction (MI) was induced by permanent ligation of the left anterior descending coronary artery (LAD) as described previously ^63^. Briefly, mice received an intraperitoneal injection of buprenorphine (0.1 mg/kg) before surgery and were then anesthetized with 2% isoflurane (in pure oxygen, 1 L/min). For post-operative analgesia, buprenorphine was administered 6 to 8 h after the procedure and again the next morning. Myocardial infarction was achieved by permanent ligation of the left anterior descending coronary artery with a 10-0 nylon surgical suture (Ethicon). The ligature was placed 1 mm below the left atria. Mice in the sham groups went through the same procedure but did not have a ligature on their artery. The surgeon was blinded to the genotype of the animals. Surviving mice were euthanized 3-or 5-days post-surgery. Cardiac structure and function were evaluated by transthoracic echocardiographic imaging on the day before the surgery or on the morning of the procedure and again prior to sacrifice.

### Determine the area at risk and infarct size

To determine the possible effects of MK2-deficiency on the area at risk (AAR) and infarct size, separate groups of wild-type and MK2-deficient mice underwent LAD ligation and were perfused with 2% Evans blue dye 30 min after ligation. Hearts were then removed, washed in 0.9% saline, and trimmed of atria and adipose tissues. Hearts were then wrapped in plastic wrap, placed in a -80 °C freezer for 5 min, cut into 4-5 transversal sections of 2 mm thickness, photographed, incubated for 20 min in 1% 2,3,5-triphenyl tetrazolium chloride (TTC) in PBS at 37 °C, fixed using 10% formalin in PBS, and photographed again to reveal the infarct area within the AAR. The AAR was the area not stained with Evans blue and was expressed as a percentage of the total left ventricular (LV) area. The infarct area (IA) was the area not stained in red by TTC and was expressed as a percentage of the AAR.

### Diagnosis of heart rupture

All non-surviving mice underwent a necropsy to identify the cause of their death. Death due to heart rupture was diagnosed as the presence of coagulated blood around the heart and in the chest cavity.

### Transthoracic echocardiography and calculations

Echocardiographic imaging was performed on mice anesthetized with 2% isoflurane (in pure oxygen, 1 mL/min) within 24 h of surgery and immediately before sacrifice. Images were acquired using a Vivid 7 Dimension system (GE Healthcare, Horten, Norway) and a i13L probe (10-14 MHz) by a technician who was blinded as to the genotype of the mice. The measurements obtained are the average of three consecutive cardiac cycles. Echocardiographic imaging of LV structure and function were as described previously ^63, 64^. Two-dimension echocardiography was used to visualize the MI. Left ventricular anterior wall thickness at end cardiac diastole (LVAWd) and LV posterior wall thickness at end cardiac diastole (LVPWd) as well as the LV dimension at end cardiac diastole (LVDd) and at end cardiac systole (LVDs) were measured using a parasternal short-axis view at the level of the papillary muscles by using M-mode echocardiography. Left atrium dimension at end cardiac systole (LADs) and diastole (LADd) were also measured using the same mode. These parameters were then used to calculate the LV mass (LVmass) using this equation: 1.055 · ((LVDd + LVAWd + LVPWd)^3^ - (LVDd)3). The parameters were also used to calculate the fractional shortening (FS) by using the following equation: (LVDd − LVDs) / LVDd · 100% ^65^. The formula available in the Vivid 7 operating system was used to calculate the LV ejection fraction (EF). LV regional contractility was assessed by using tissue Doppler imaging (TDI) to determine basal lateral wall systolic contractile velocity (SL) and basal septum systolic contractile velocity (SS). Pulsed wave Doppler was used in apical four-chamber view to obtain various LV diastolic parameters such as the trans mitral flow (TMF), the early filling deceleration time (EDT), the early filling deceleration rate (EDR), the mitral valve closure to opening time (MVCO), the trans-mitral early (E) and late atrial (A) filling velocities. Pulsed-wave Doppler was used to obtain the LV ejection time (LVET) by measuring the time starting from the beginning to the end of the LV outflow. These measurements were used to calculate the global myocardial performance index (global MPI) by using the following formula: (MVCO − LVET) / LVET · 100%. The systolic flow of pulmonary venous flow (PVF) (S) and the diastolic flow of PVF (D) were both also obtained using the pulsed wave Doppler. The mitral annulus was viewed by TDI to measure velocities during early filling on the septal segment (septal Em) and lateral segment (lateral Em) as well as the atrial filling for the septal (septal Am) and the lateral (lateral Am) segments. These parameters were used to calculate values for lateral and septal E/Em. The images taken by pulsed-wave Doppler in apical five-chamber view were used to determine the isovolumetric relaxation time (IVRT). As MK2-deficient mice are bradycardic, the heart rate-corrected IVRT was calculated as follows: corrected IVRT (IVRTc) = IVRT / RR1^/^2. The RR interval was acquired using simultaneously recorded ECG. The septal and lateral MPI were calculated using the following equation (b − a) / a · 100%. The value identified as “a” is the interval of time from the start to the end of the SL or SS for lateral and septal MPIs, respectively. Similarly, “b” corresponds to the interval of time from the end of Am to the beginning of Em measured at the lateral and septal annular segments for lateral and septal MPIs, respectively. LV wall motion was scored using a scale numbered from 1 to 5: 1 = normal, 2 = hypokinesis, 3 = akinesis, 4 = dyskinesis and 5 = aneurysmal. The wall motion score index (WMSI) was calculated by using the following equation: sum of all scores / number of segments viewed. Right ventricular (RV) diameter at end diastole (RVDd), RV anterior wall thickness at end diastole (RVAWd) and tricuspid annulus plane systolic excursion (TAPSE) were measured using M-mode echocardiography. Pulsed wave Doppler was used to obtain the acceleration time (AT) and RV ejection time (RVET) of the pulmonary artery flow. The same method was used to evaluate trans-tricuspid flow early filling velocity (Et), late (atrial) filling velocity (At), early filling deceleration rate (EtD rate), early filling deceleration time (EtDT), and tricuspid valve closure to opening time (TVCO). Tissue Doppler imaging of the tricuspid annulus was used to measure the right ventricular lateral wall peak systolic velocity (SR), lateral tricuspid annulus peak velocity during early filling (Em), and lateral tricuspid annulus peak velocity during late (atrial) filling (Am). The RV MPI was calculated in the same manner as that of the LV.

### Histological analysis

Mice were anesthetized with 2% isoflurane (in pure oxygen, 1 L/min), hearts removed, and perfused with a saline solution (0.9% sodium chloride and heparin 2 USP units/mL) to clear the blood in the arteries, followed by a second perfusion with 10% formalin. A transversal cut was then made between the ligature and the apex of the heart, and the two halves were then placed in a cassette and immersed in 10% formalin solution. Staining and collagen quantification were performed in the histology facility in the laboratory of Dr. Martin G. Sirois at the Montreal Heart Institute by a histology technician blinded to the genotype of the sample. Following a 24 h fixation step in 10% formalin, hearts were dehydrated by immersion in solutions of increasing alcohol concentrations (70%, 95%, 100%), followed by xylene, embedded in paraffin, cut in 6 µm transversal sections, and mounted on charged slides. Sections from all mice in the study underwent histochemical and immunohistochemical assessment. In preparation for immunohistochemistry, tissue sections were treated with citrate buffer (pH 6.0) for antigen retrieval and 3% hydrogen peroxide solution to block endogenous peroxidase activities prior to blocking for 20 min in phosphate buffered saline (PBS) containing 10% serum (same species as secondary antibody). Sections were then incubated with primary antibody diluted in PBS containing 1% normal serum overnight in a humidified chamber at 4 °C. The primary antibodies used were a mouse monoclonal antibody against smooth muscle *α*-actin (*α*-SMA, a myofibroblast marker; A-2547, Sigma-Aldrich) and rabbit polyclonal antibodies against myeloperoxidase (MPO, a neutrophil and monocyte marker; ab65871, Abcam), CD206/mannose receptor (a macrophage marker; ab64693, Abcam), and CD31/PECAM-1 (an endothelial cell marker; sc-1506, Santa Cruz Biotechnology). The negative control groups were incubated in the absence of primary antibody. After washing, tissue sections were incubated with a biotin-conjugated secondary antibody for 30 min, washed again, and incubated with horseradish peroxidase-conjugated streptavidin (Vector Labs) for 30 min. Immunoreactivity was visualized using the chromogenic peroxidase substrate 3,3’-diaminobenzidine (Vector Labs). Finally, sections were counter-stained using hematoxylin and mounted using Permount. Images of the heart sections were taken at 4X and 20X using an Olympus BX46 microscope. Analysis of the infarct area as well as collagen quantification in the infarct area were performed using Image Pro Plus software version 7.0 (Media Cybernetics, Silver Spring, MD) and expressed as a percentage of the total tissue area of each section. Images of MPO-immunoreactivity were acquired at 20X magnification whereas those of CD206, CD31, or *α*-SMA immunoreactivity were acquired at 40X using an Olympus BX46 microscope.

### Isolation and culture of cardiac ventricular fibroblasts

Fibroblasts were isolated from male MK2^+/+^ and MK2^-/-^ mice aged between 11 to 13 weeks as described previously ^63, 64, 66^. Mice were sacrificed by exsanguination following a pentobarbital injection. Hearts were removed and immediately immersed in sterile PBS (137 mM NaCl, 2.7 mM KCl, 4.2 mM Na2HPO4·H2O, pH 7.4) at 37 °C. After removing the atria and adipose tissue, the ventricles were cut into pieces of approximately 1 mm^2^, which were then subjected to a series of digestions in a dissociation medium (116.4 mM NaCl, 23.4 mM HEPES, 0.94 mM NaH2PO4·H2O, 5.4 mM KCl, 5.5 mM dextrose, 0.4 mM MgSO4, 1 mM CaCl2, 1 mg/mL BSA, 0.5 mg/mL collagenase type IA, 1 mg/mL trypsin, 0.020 mg/mL pancreatin, pH 7.4). with gentle shaking on an orbital mixer placed in a humidified incubator at 37 °C under a 5% CO2 atmosphere. After 10 minutes, the supernatant, containing the cells released from the tissue, were collected, and centrifuged at 1500 rpm for 5 min. After 10 cycles of digestion and centrifugation, the cell pellets were resuspended in 4 mL of Medium 199 (M199, Sigma-Aldrich) supplemented with 10% FBS, 2% antibiotics (streptomycin and penicillin, Hyclone) and 2% amphotericin B (Gibco), plated onto two 35 mm petri dishes, and incubated in a humidified incubator at 37 °C in a 5% CO2 atmosphere. The media was changed after 150 minutes, to remove cell debris, and again the following morning. These cells were referred to as passage 0.

### Scratch-wound assays

Fibroblast motility was assessed by scratch-wound assay in passage 0 fibroblasts cultures. Fibroblasts were seeded directly in 12 well plates and the media was changed every 24 h until cultures reached 80% confluence. Cells were then washed with PBS and maintained in serum-free M199 for 18 h. A scratch was created across the center of each well with a P1000 pipette tip and the plates were washed with PBS to remove the debris. Different treatments were used: serum-free media, media containing serum, and media containing serum and angiotensin II. Images were acquired immediately after creating the scratch (time zero) and again 24 h later using a Nikon ELWD 0.3/OC 75 camera (magnification of 4X). ImageJ version 1.48 was used to assess the percentage of the open wound remaining after 24 h.

### RNA extraction and RT^2^ Profiler PCR Arrays

The LV tissue from infarcted hearts was separated into healthy and infarcted tissue. The LV from sham mice was left intact. Tissue samples were then submerged in 2-methylbutane, snap frozen with liquid nitrogen, and stored frozen at -80 °C until further analysis. In preparation for RNA extraction, frozen tissue was pulverized to a fine powder under liquid nitrogen using a mortar and pestle. Powdered tissue was combined with 500 µL of TRIzol reagent and homogenized with a single 10 s burst using a Polytron PT 2100 homogenizer. Total RNA was then purified using RNeasy Mini Kits (QIAGEN Inc.) according to the manufacturer’s instructions. The concentration and purity of each RNA sample were evaluated using a NanoDrop ND-1000 spectrophotometer and only the samples with a 260/280 absorbance ratio above 1.8 were used. First strand cDNA was synthesized from 0.5 mg of total RNA using QIAGEN RT2 First Strand Kits and transcripts for mouse cytokines and chemokines were quantified using qPCR microarrays comprising 96-well plates precoated with primers (QIAGEN PAMM-150Z). Each 96-well plate also contained primers for 5 housekeeping genes as well as positive and negative controls. qPCR was performed using an ABI StepOnePlus instrument according to the instructions provided by the manufacturer. Data was analyzed according to the instructions provided in the RT^2^ Profiler PCR Array Data Analysis v3.5 Handbook using the software on the QIAGEN website (https://dataanalysis.sabiosciences.com/pcr/arrayanalysis.php) and normalized to internal controls. Changes were considered significant when *P* < 0.05.

### Statistical analysis

Normality tests (Shapiro-Wilk) were performed on all data. The Mandel-Cox test was used to analyze survival curves. Data are presented as mean ± SEM or median (1^st^ quartile, 3^rd^ quartile). For multiple comparisons of means involving a combination of 2 independent factors such as surgery (LAD ligation, sham) or genotype (MK2^+/+^, MK2^-/-^), 2-way ANOVA followed by Tukey’s post hoc tests were performed. When a lognormal distribution was indicated, data was log-transformed before conducting the ANOVA. All tests were two-sided. A *P* value < 0.05 was considered significant. Statistical analyses were performed using Prism version 9.4.1 for Mac OS X (GraphPad Software, La Jolla, CA).

## Results

### MK2-deficiency decreased mortality after LAD ligation

Through phosphorylation of RNA-binding proteins, MK2 regulates the stability of several cytokine transcripts ^43, 46^. As the first phase of wound-healing post-MI involves an inflammatory response, we assessed the effects of an MK2-deficiency on the inflammatory phase of post-MI wound healing. Myocardial infarctions were induced by permanent ligation of the left anterior descending coronary artery and mice were sacrificed on the third or fifth day following the surgery, which roughly corresponds to the middle and the end of the inflammatory response. All sham-operated (MK2^+/+^: 39/39; MK2^-/-^: 38/38) and ligated mice (MK2^+/+^: n = 45/45; MK2^-/-^: n = 41/41) survived to day 3 post-MI (**Figure 1A**). However, 5 days post-MI, survival of MK2^-/-^ mice was significantly greater (100%, n = 21/21) than MK2^+/+^ mice (79%, n = 23/29, Mandel-Cox test: *P* = 0.0285) (**Figure 1B**). Post-mortem examinations revealed that all deaths in MI-MK2^+/+^ mice occurring on days 3 - 5 were due to heart rupture, as determined by both the accumulation of blood in the chest cavity and the presence of a clot on the heart ^67^. In addition, one MI-MK2^+/+^ mouse had to be sacrificed on day 4 as it was in distress. A similar number of MK2^+/+^ and MK2^-/-^ mice did not survive the first hour after the surgery (MK2^+/+^: n = 3; MK2^-/-^: n = 2). These 6 mice were excluded from the study. Hence, mortality was reduced in MK2-deficient mice during the inflammatory phase of post-MI wound healing.

**Figure 1.**
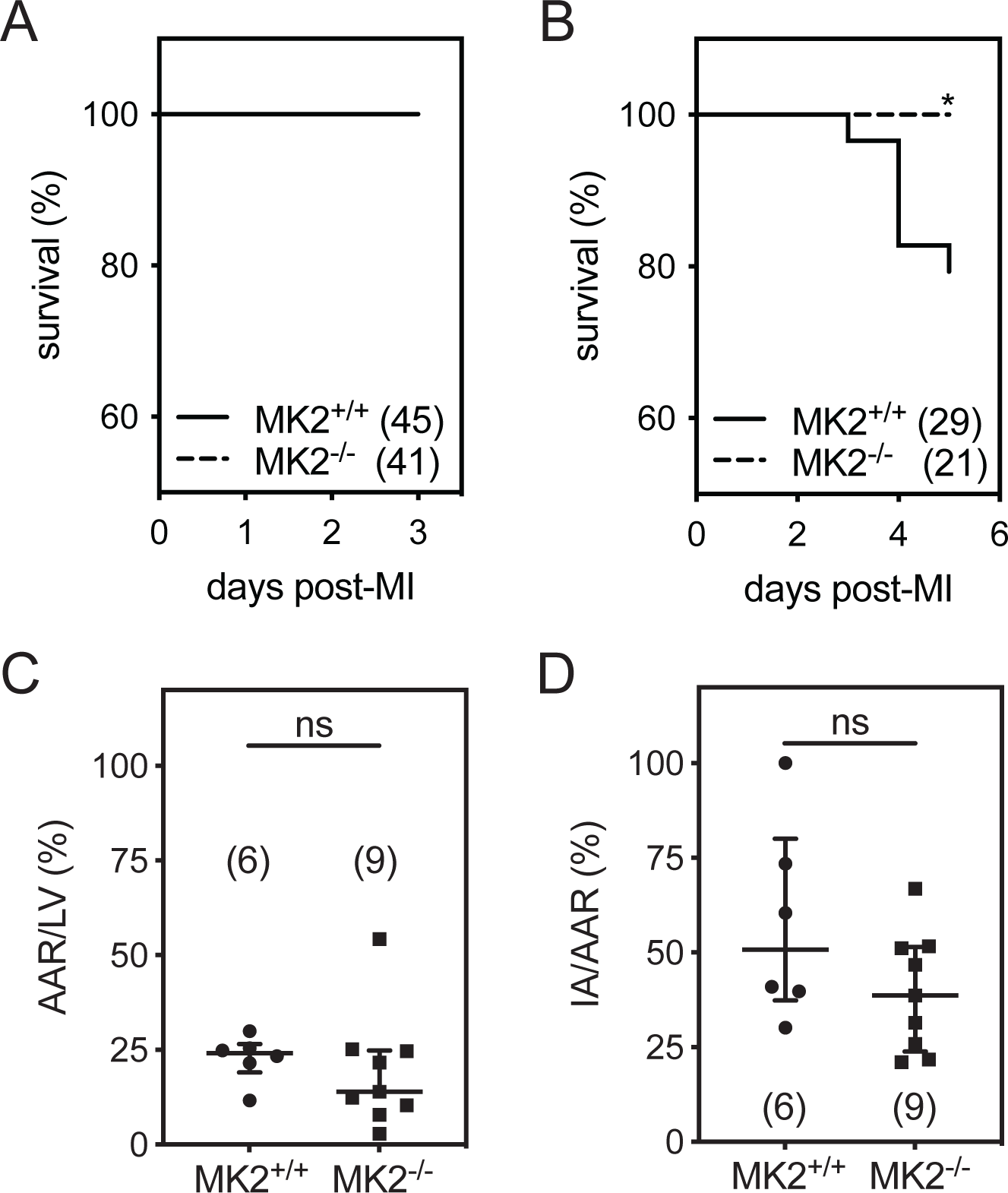
MK2-deficiency improved post-MI survival during the inflammatory phase and did not alter area at risk or infarct size. **A**, Kaplan-Meier survival curve for MK2^+/+^ (solid line, n = 45) and MK2^-/-^ (dashed line, n = 41) mice in the first 3 days post-MI. **B**, Kaplan-Meier survival curve for MK2^+/+^ (solid line, n = 29) and MK2^-/-^ (dashed line, n = 21) mice in the first 5 days post-MI (*, Mandel-Cox test: *P* = 0.0285). **C**, Area at risk (AAR) was assessed by infusion of Evans blue dye in MK2^+/+^ (n = 6) and MK2^-/-^ mice (n = 9) 30 minutes after ligation of the left anterior descending coronary artery. The AAR, which corresponds to the area in which the dye was excluded, is expressed as a percentage of the left ventricular (LV) wall area. **D**, Infarct area (IA) was assessed by 2,3,5-triphenyl tetrazolium chloride (TTC) staining in MK2^+/+^ (n = 6) and MK2^-/-^ mice (n = 9). Non-viable myocardium, which is not stained by TTC, is expressed as a percentage of the AAR. The number of animals in each group is indicated in parentheses. Shapiro-Wilk tests for normality were performed on all data. Results shown are median (first quartile and third quartile). Mann-Whitney tests were performed for statistical comparisons.

### MK2-deficiency did not alter the area at risk or infarct size

We examined the possible effect of an MK2-deficiency on the size of the area at risk (AAR) following LAD ligation by infusing a group of MK2^+/+^ and MK2^-/-^ mice with Evans blue dye, which is excluded from the AAR, 30 min after ligation. MK2-deficiency had no effect on the size of the AAR [expressed as a percentage of left ventricular (LV) area (MK2^+/+^: 22.8 ± 2.5%, n = 6; MK2^-/-^: 19.2 ± 5.1%, n = 9; *P* > 0.05: **Figure 1C**)]. Similarly, 2,3,5-triphenyl tetrazolium chloride (TTC), which stains viable myocardium, revealed that, after 30 min of ligation, infarcts were of similar size in MK2^+/+^ and MK2^-/-^ mice [expressed as a percentage of AAR (MK2^+/+^: 57.5 ± 10.7%, n = 6; MK2^-/-^: 39.5 ± 5.3%, n = 9; *P* > 0.05: **Figure 1D**)].

### MK2-deficiency attenuated LV dilation post-MI

The effects of MK2-deficiency on LV and RV structural and functional remodeling post-MI were assessed by echocardiographic imaging (**Tables 1 and 2**). Generally speaking, three- and five-days post-MI, certain genotype-specific differences were detected in the structural or functional alterations resulting from MI. Both MK2^+/+^ and MK2^-/-^ mice displayed reduced LV (ejection fraction) and RV (tricuspid annular plane systolic excursion) systolic function, left atrial (LA) dilation, increased wall motion score index (WMSI), and increased (indicating poorer performance) in LV and RV myocardial performance indices. Tissue Doppler imaging revealed little genotype-dependent effect on the MI-mediated changes in mitral annulus velocity: the exception being the peak systolic velocity of the septal segment (SS), which was reduced in MI-MK2^-/-^ mice, relative to MI-MK2^+/+^ mice, whereas the lateral segment velocity (SL) was not significantly affected by the absence of MK2 (**Figure 2A and 2B**). Note that in comparing structure, 12-week-old MK2-deficient mice were smaller than age-matched littermates (**Table 1**) ^60^. However, 2-way ANOVA also revealed significant genotype x surgery interactions post-MI. The thickness of the LV posterior wall (LVPWd) was significantly increased in MI-MK2^-/-^ but not MI-MK2^+/+^ mice 5-d post-MI. Left ventricular internal diameter at end-diastole (LVDd) and end-systole (LVDs) were significantly smaller in MI-MK2^-/-^ than MI-MK2^+/+^ mice 5-d post-MI (**Table 1, Figure 2C and 2D**) as was LV volume at end-diastole (LVVd) (**Figure 2E**). Although the LV volume at end-systole (LVVs) was also significantly smaller in MI-MK2^-/-^ than MI-MK2^+/+^ mice 5-d post-MI (**Table 1, Figure 2F**), no interaction effect was detected (*P* > 0.05). Examining transmitral flow velocities during diastole showed early diastolic filling velocity (E) and the ratio of early to late diastolic filling velocities (E/A) were greater in MI-MK2^-/-^ mice than MI-MK2^+/+^ mice 3-d post-MI as was the peak upper pulmonary venous flow during diastole (upper D) (surgery x genotype interaction: *P* < 0.05). The deceleration time and deceleration rate of the early wave of diastolic filling (EDT, ED rate) as well as heart rate-corrected isovolumetric relaxation time (IVRTc) in MI-MK2^-/-^ and MI-MK2^+/+^ mice did not differ significantly. Similarly, the ratio of peak E wave velocity to mitral annulus peak velocity during early diastolic filling (E/Em), an indicator of LV filling pressure, in MI-MK2^-/-^ and MI-MK2^+/+^ mice, did not differ significantly. These data suggest that the deficiency of MK2 attenuated LV dilation post-MI but was neither beneficial nor detrimental to cardiac systolic or diastolic function.

**Figure 2.**
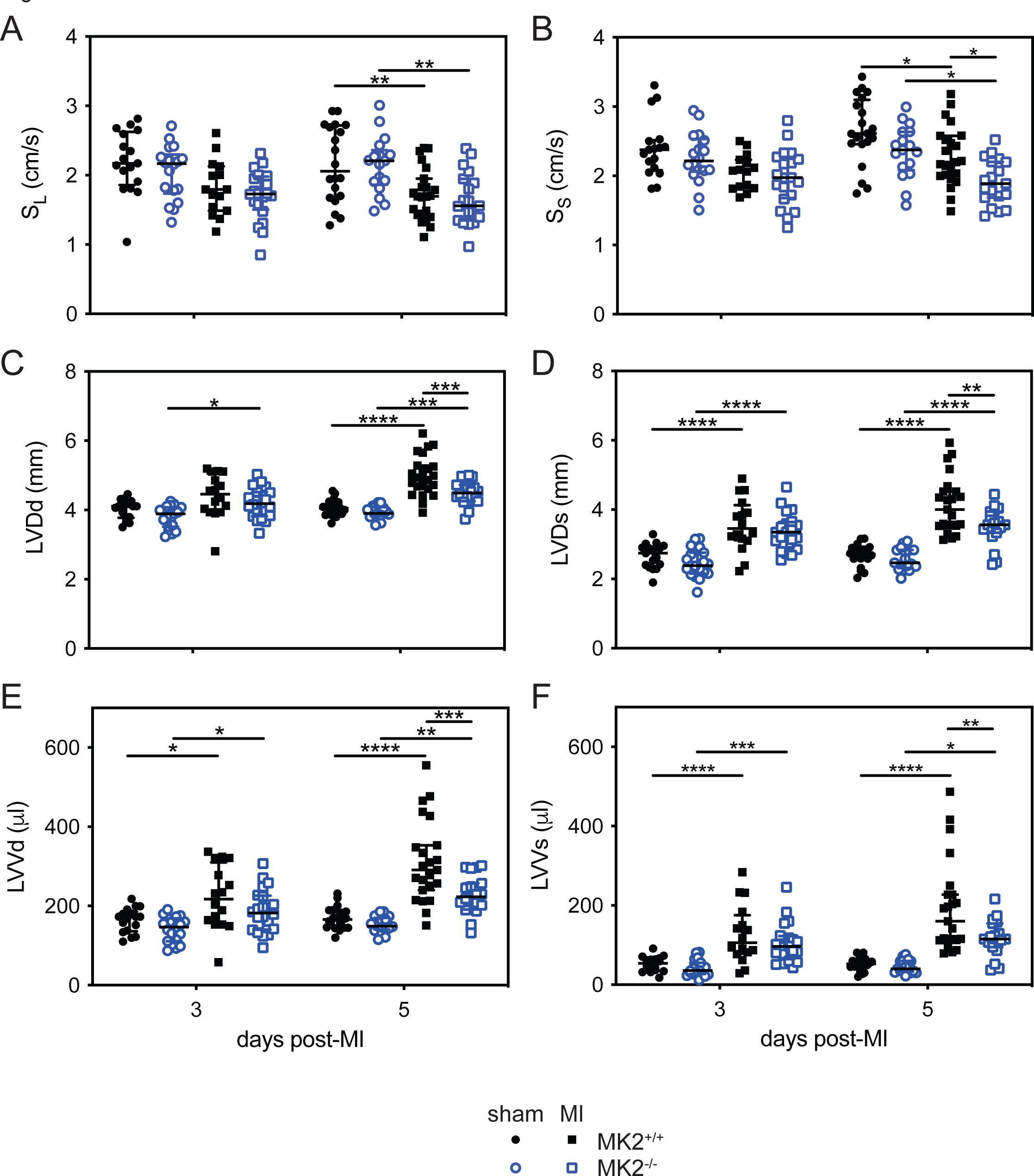
MK2-deficiency attenuated LV dilation five days post-MI. **A**, Basal lateral systolic contractile velocity (SL). **B**, Basal septal systolic contractile velocity (SS). **C**, Left ventricular (LV) dimension at end cardiac diastole (LVDd). **D**, LV diameter at end cardiac systole (LVDs). **E**, LV volume at end cardiac diastole (LVVd). **F**, LV volume at end cardiac systole (LVVs). Sham-MK2^+/+^ (solid black circles, 3 days n = 17, 5 days n = 20), sham-MK2^-/-^ (open blue circles, 3 days n = 18 - 19, 5 days n = 17), MI-MK2^+/+^ (solid black squares, 3 days n = 15 - 16, 5 days n = 20 -23) and MI-MK2^-/-^ (open blue squares, 3 days n = 20, 5 days n = 19) mice. Data are presented as median (first quartile and third quartile). Shapiro-Wilk tests for normality were performed on all data. Data with a lognormal distribution were log-transformed prior to statistical comparison by two-way ANOVA, which included a factor for surgery (sham, LADL), a factor for genotype (MK2^+/+^, MK2^-/-^), and a surgery x genotype interaction term. The ANOVA was followed by Tukey’s post hoc tests for multiple comparisons. \**P* < 0.05, \*\**P* < 0.01, \*\*\**P* < 0.001, \*\*\*\**P* < 0.0001.

The absence of MK2 also altered remodeling of the RV post-MI. Although the group means for right ventricular internal diameter at end-diastole (RVDd) did not differ significantly, there was a significant interaction effect 3-d post MI (*P* < 0.05): RVDd increased in MI-MK2^+/+^ mice, relative to sham-MK2^+/+^ mice, whereas it decreased in MI-MK2^-/-^ mice (**Table 2**). The absence of MK2 also altered the effect of MI on transtricuspid flow early filling velocity (Et) 3-d post-MI. In MI-MK2^+/+^ hearts, Et was increased, relative to sham mice, whereas it was decreased in MI-MK2^-/-^ hearts. A similar effect was observed on the ratio of Et to tricuspid annulus peak velocity during early diastolic filling (Et/Em). An interaction effect (*P* < 0.05) was observed for tricuspid Em 5-d post-MI, which was actually a result of Em in MI-MK2^+/+^ hearts decreasing, relative to sham-MK2^+/+^, to peak velocities observed in sham-MK2^-/-^ and MI-MK2^-/-^ mice. Tricuspid Em was slower in sham-MK2^-/-^ hearts and did not undergo further reduction as a result of MI.

**Table 2.**
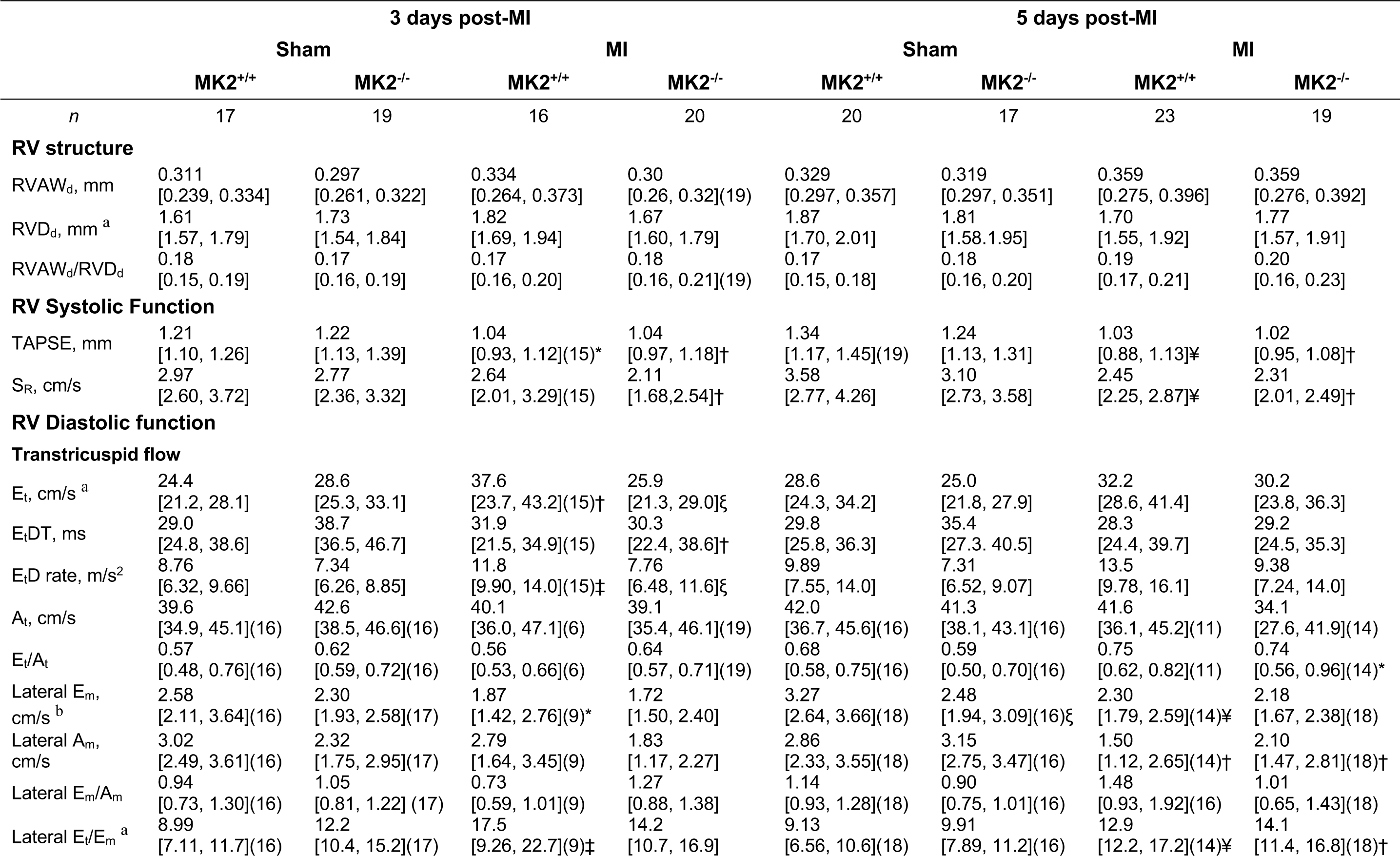
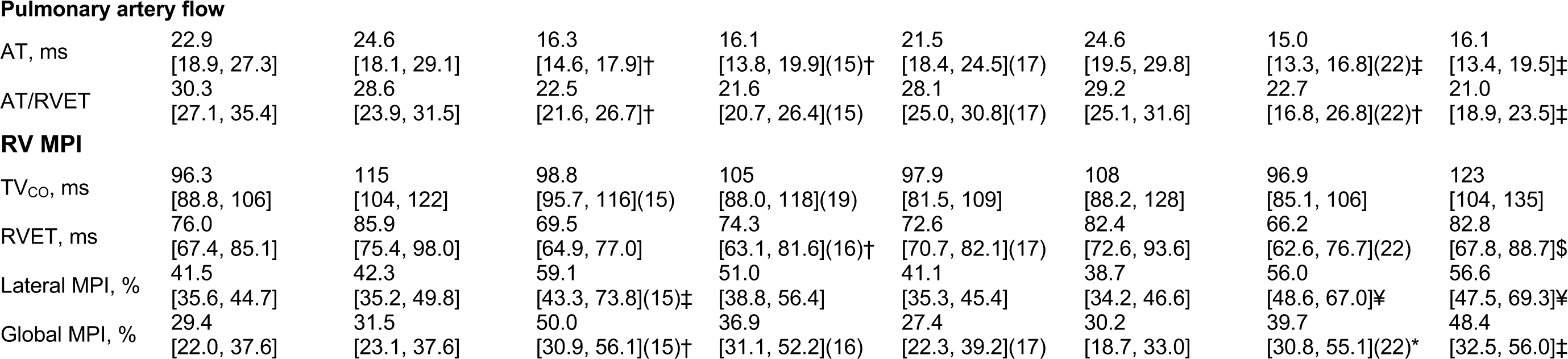
Echocardiographic parameters of RV structure and function in 12-week-old MK2^+/+^ and MK2^-/-^ mice at 3 or 5 days post-MI. Data are reported as median [1^st^ quartile, 3^rd^ quartile]. The *n* numbers are as shown in the first row of the table unless otherwise indicated in parentheses. Am, tricuspid annulus peak velocity during late (atrial) diastolic filing; AT, pulmonary arterial flow acceleration time; At, transtricuspid flow late (atrial) filling velocity; Em, tricuspid annulus peak velocity during early diastolic filing; Et, transtricuspid flow early filling velocity; EtD rate, transtricuspid early filling deceleration rate; EtDT, transtricuspid early filling deceleration time; MK2, mitogen-activated protein kinase–activated protein kinase-2; MPI, myocardial performance index; RV, right ventricular; RVAWd, right ventricular anterior wall thickness at end cardiac diastole; RVDd, right ventricular diameter at end cardiac diastole; RVET, right ventricular ejection time; SR, right ventricular lateral wall systolic velocity; TAPSE, tricuspid annulus plane systolic excursion; and TVCO; tricuspid valve closure to opening time. Shapiro-Wilk tests for normality were performed for all data: if a lognormal distribution was indicated, data was log-transformed prior to statistical comparison by two-way ANOVA, including a factor for surgery (sham, MI), a factor for genotype (MK2^+/+^, MK2^-/-^), and a surgery x genotype interaction term. The ANOVA was followed by Tukey’s post hoc tests for multiple comparisons to compare means. *: p < 0.05 vs Sham. †: p < 0.01 vs Sham. ‡: p < 0.001 vs sham. ¥: p < 0.0001 vs Sham. $: p < 0.05 vs MK2^+/+^. ξ: p < 0.01 vs MK2^+/+^. §: p < 0.001 vs MK2^+/+^. ^+/+^. ^a^: *P* < 0.05 for surgery x genotype interaction 3-days post-MI. ^b^: *P* < 0.05 for surgery x genotype interaction 5-days post-MI.

### MK2-deficiency did not alter infarct area size or collagen content 3- and 5-days post-MI

Reparative fibrosis post-MI involves the deposition of collagen to provide structural support to the LV wall ^3^. Impaired collagen deposition can lead to thinning of the LV wall whereas excessive deposition can cause LV stiffness: both events can negatively affect heart function ^3, 5^. The larger a scar area becomes, the more severely function is impaired ^3, 5^. As a result, infarct size and collagen content are crucial determinants of the long-term outcomes post-MI. For histological analysis, hearts were first cut along the short axis, as shown in Figure 3A, resulting in upper (A) and lower regions (B) of the infarct (**Figure 3A**). The upper sections allowed a better assessment of the scar border, whereas the lower sections permitted a better assessment of scar morphology. Sections from both ‘A’ and ‘B’ were stained using Masson’s trichrome, which colors healthy tissue red and collagen blue. MK2-deficiency did not alter the infarct area (**Figure 3B and 3C**) 3- or 5-days post-MI in either the upper (section A) or lower (section B) regions of the LV. As one would expect this early in the wound-repair process, the collagen content in the infarct area (**Figure 4**) was low 3 days post-MI and had increased slightly 5 days post-MI. In addition, 5 days post-MI, the collagen content in the infarct region of the ‘A’ sections was less in MI-MK2^-/-^ than MI-MK2^+/+^ mice (**Figure 4B**).

**Figure 3.**
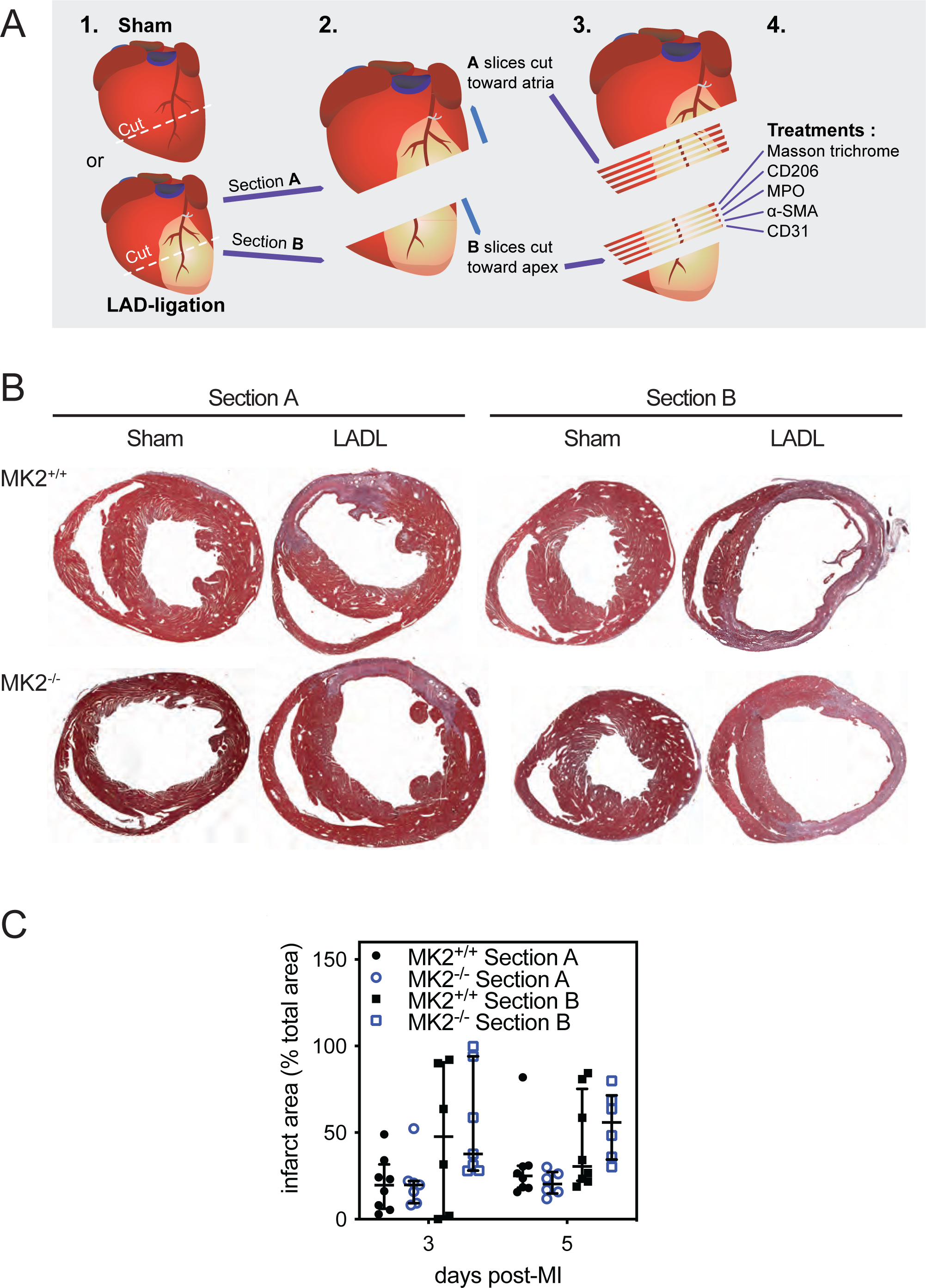
MK2-deficiency did not alter infarct area 3- or 5-days post-MI. **A**, Representative images of a hearts cut along the short axis (1). Sections taken from the upper region of the infarct are identified as “Section A” and those from the lower region as “Section B” (2, 3). Sections from both regions were employed for histological (Masson’s trichrome stain) or immunohistochemical stain (4). **B**, Representative Masson’s trichrome stained short-axis sections of the ventricular myocardium from sham and ligated MK2^+/+^ and MK2^-/-^ mice 5 days post-myocardial infarction (MI) showing collagen deposition (blue) and healthy tissue (red). **C**, Infarct area, identified by Masson’s trichrome staining, expressed as a percentage of the total myocardial area from MK2^+/+^ (solid black circles: Section A, n = 8; solid black squares: Section B, n = 6 - 8) and MK2^-/-^ (open blue circles: Section A, n = 6 - 7; open blue squares: Section B, n = 6 - 7) mice. Data are presented as median (first quartile and third quartile). Shapiro-Wilk tests for normality were performed on all data. Mann-Whitney tests were performed for statistical comparisons between MK2^+/+^ and MK2^-/-^.

**Figure 4.**
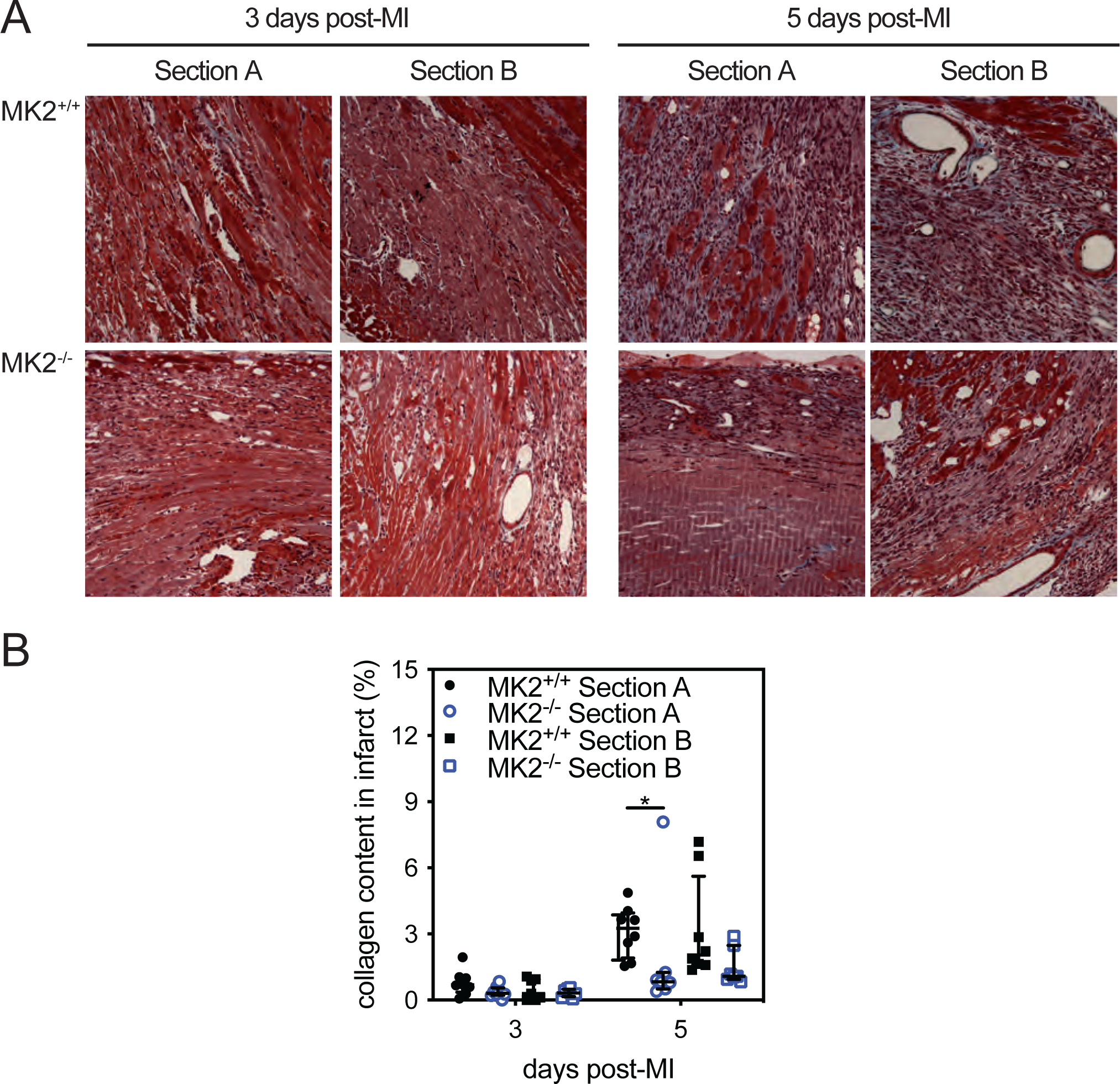
MK2-deficiency did not hinder collagen deposition 3- and 5-days post-MI. **A**, Masson’s trichrome stained short-axis sections of the ventricular myocardium from MK2^+/+^ and MK2^-/-^ mice 3- and 5-days post-myocardial infarction (MI) showing collagen deposition (blue) and healthy tissue (red). Hearts were cut along the short axis through the center of the infarct to yield upper (Section A) and lower, (Section B) regions. **B**, Collagen content expressed as a percentage of the total infarct area from MK2^+/+^ (solid black circles: Section A, n = 8; solid black squares: Section B, n = 7 and 8) and MK2^-/-^ (open blue circles: Section A, n = 7 - 8; open blue squares: Section B, n = 7 - 8) mice. Images were analyzed by color segmentation using Image Pro Plus version 7.0 (Media Cybernetics, Silver Spring, MD). Shapiro-Wilk tests for normality were performed on all data. Data are presented as median (first quartile and third quartile). Mann-Whitney tests were performed for statistical comparisons between MK2^+/+^ and MK2^-/-^. \**P* < 0.05.

### MK2-deficiency did not affect neutrophil or macrophage recruitment to the infarct or peri-infarct regions

As the inflammatory phase of wound repair is essential for appropriate healing and the absence of MK2 did not result in increased mortality within the first 5-days post-MI, we next sought to characterize the effects of MK2-deficiency on the inflammatory response in infarcted hearts. Neutrophils and monocytes/M1 macrophages, the first responders to the inflammatory factors released by necrotic myocytes ^20^, begin to clear the debris caused by the ischemia and participate in the inflammatory response by secreting numerous cytokines ^27, 64, 68^. To determine if the absence of MK2 affected the recruitment of neutrophils or monocytes to the infarct or peri-infarct regions, tissue sections were decorated with antibodies against myeloperoxidase (MPO). MPO immunoreactivity was sparse in the myocardium of sham hearts and the non-infarcted myocardium hearts from both MI-MK2^+/+^ and MI-MK2^-/-^ mice. The abundance of MPO immunoreactivity was greater in the peri-infarct and infarct regions, relative to both sham hearts and the non-infarcted myocardium of infarcted hearts; however, MPO immunoreactivity did not differ significantly between MK2^+/+^ and MK2^-/-^ mice 3- or 5-days post-MI (**Figure 5**).

**Figure 5.**
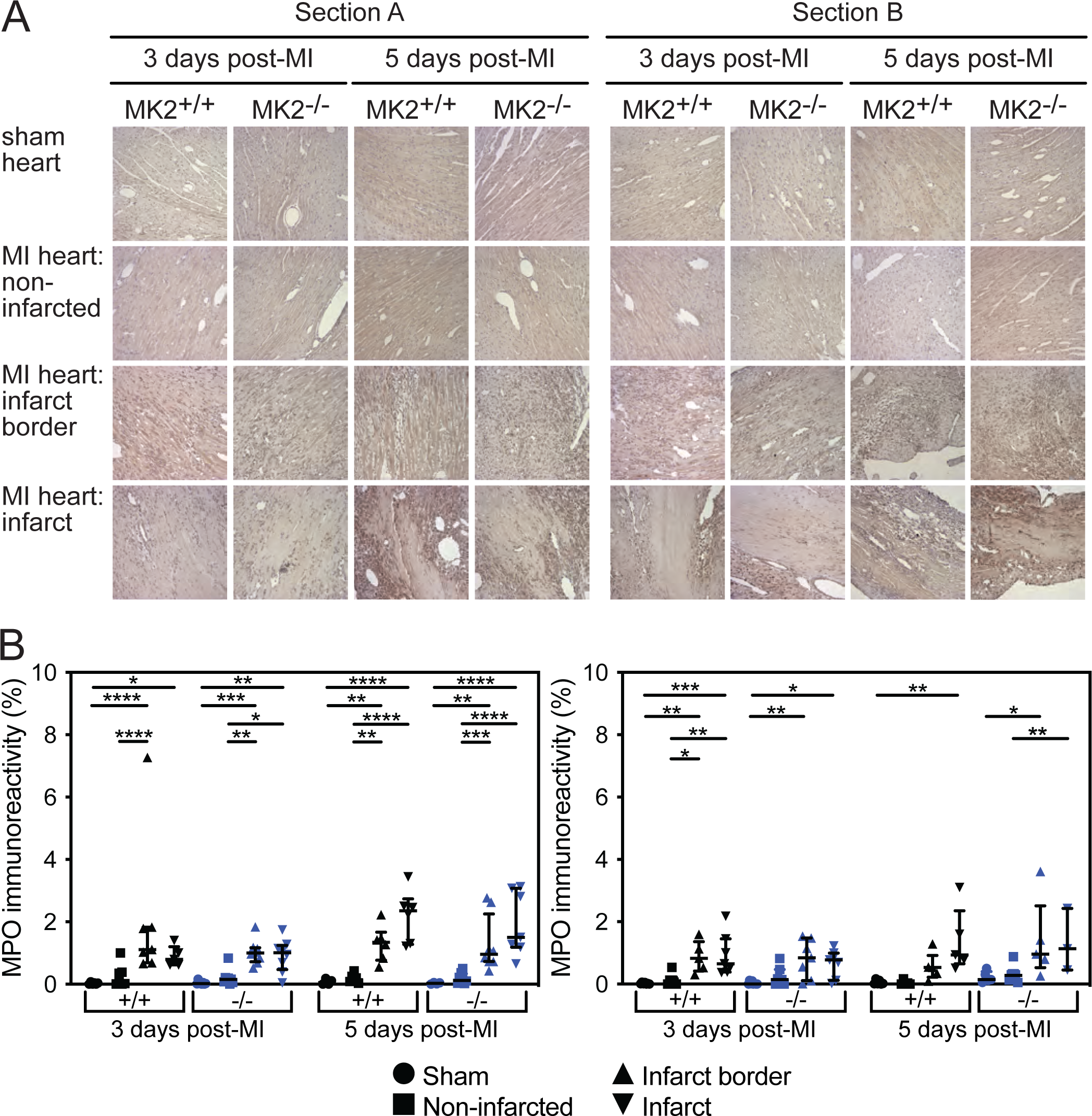
MK2-deficiency did not hinder recruitment of neutrophils and monocytes to the peri-infarct and infarct areas 3- and 5-days post-MI. A. Representative images of immunohistochemical staining for myeloperoxidase (MPO, dark brown), a neutrophil and monocyte marker, in sham and infarcted hearts from MK2^+/+^ and MK2^-/-^ mice euthanized 3- and 5-days post-MI. Hearts were cut along the short axis through the center of the infarct to yield upper (Section A) and lower, (Section B) regions. B, MPO immunoreactivity expressed as a percentage of the total field area from sham-MK2^+/+^ (black circles, n = 4 - 7), MI-MK2^+/+^ (non-infarcted tissue: black squares, n = 6 - 8; peri-infarct: black triangles, n = 5 - 7; infarct: inverted black triangles, n = 5 - 7), sham-MK2^-/-^ (blue circles, n = 4 - 8), and MI-MK2^-/-^ (non-infarcted tissue: blue squares, n = 6 - 8; peri-infarct: blue triangles, n = 6 - 8; infarct: inverted blue triangles, n = 3 - 8) hearts. Images were analyzed by color segmentation using Image Pro Plus version 7.0 (Media Cybernetics, Silver Spring, MD). Shapiro-Wilk tests for normality were performed on all data. Data are presented as median (first quartile and third quartile). Data with a lognormal distribution were log-transformed prior to statistical comparison by two-way ANOVA, which included a factor for surgery (sham, MI), a factor for genotype (MK2^+/+^, MK2^-/-^), and a surgery x genotype interaction term. No interaction was detected (*P* > 0.05). The ANOVA was followed by Tukey’s post hoc tests for multiple comparisons. \**P* < 0.05, \*\**P* < 0.01, \*\*\**P* < 0.001, \*\*\*\**P* < 0.0001.

As the inflammatory response evolves, M1 macrophages shift to their M2 phenotype, which are then involved in resolving the inflammation and initiating tissue repair. The abundance of M2 macrophages, assessed by decorating tissue sections with anti-CD206 antibodies, was low in the myocardium of sham hearts and the non-infarcted myocardium of hearts from both MI-MK2^+/+^ and MI-MK2^-/-^ mice and greatest in the peri-infarct zone at both 3- and 5-days post-MI (**Figure 6**). Interestingly, 5-days post-MI, in the ‘A’ sections, the abundance of CD206 immunoreactivity in the peri-infarct region of MI-MK2^+/+^ hearts showed a trend towards being greater than in MI-MK2^-/-^ hearts, but the difference did not reach significance. No genotype-dependent differences were observed in the ‘B’ sections.

**Figure 6.**
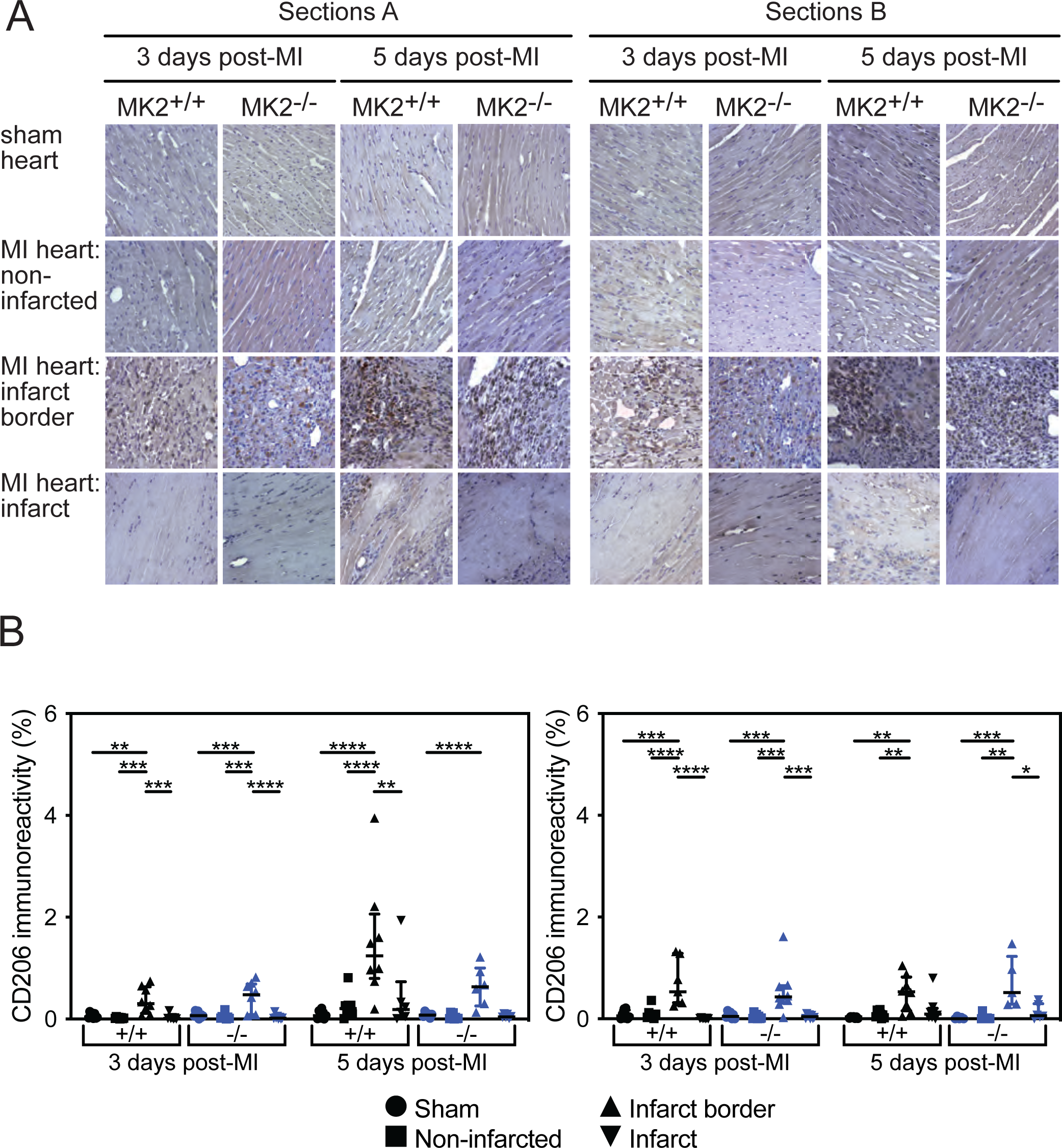
MK2-deficiency did not hinder M2 macrophage recruitment to the peri-infarct area 3- and 5-days post-MI. **A**, Representative images of immunohistochemical staining of the mannose receptor cluster of differentiation 206 (CD206, dark brown), an M2 macrophage marker, in sham and infarcted hearts from MK2^+/+^ and MK2^-/-^ mice sacrificed 3- and 5-days post-MI. Hearts were cut along the short axis through the center of the infarct to yield two pieces (Section A, upper region; Section B, lower region). **B**, CD206 immunostaining expressed as a percentage of the total field area from sham-MK2^+/+^ (black circles, n = 7 - 8), MI-MK2^+/+^ (non-infarcted tissue: black squares, n = 7 - 8; peri-infarct: black triangles, n = 7 - 8; infarct: black inverted triangles, n = 6- 8), sham-MK2^-/-^ (blue circles, n= 8), and MI-MK2^-/-^ (non-infarcted tissue: blue squares, n = 5 - 8; peri-infarct: blue triangles, n = 5 - 8; infarct: blue inverted triangles, n = 6 - 8) hearts. Images were analyzed by color segmentation using Image Pro Plus version 7.0 (Media Cybernetics, Silver Spring, MD). Shapiro-Wilk tests for normality were performed on all data. Data are presented as median (first quartile and third quartile). Data with a lognormal distribution were log-transformed prior to statistical comparison by two-way ANOVA, which included a factor for surgery (sham, MI), a factor for genotype (MK2^+/+^, MK2^-/-^), and a surgery x genotype interaction term. No interaction was detected (*P* > 0.05). The ANOVA was followed by Tukey’s post hoc tests for multiple comparisons. \**P* < 0.05, \*\**P* < 0.01, \*\*\**P* < 0.001, \*\*\*\**P* < 0.0001.

### MK2-deficiency did not alter the accumulation or distribution of myofibroblasts

The healing process following an MI involves formation of a collagen scar produced by fibroblasts that migrate into the affected area and become activated to myofibroblasts ^5, 11^. A hallmark of myofibroblasts is the expression of α-SMA ^26, 69^. Thus, α-SMA-specific antibodies were used to assess the abundance of myofibroblasts in transverse sections of infarcted and sham hearts (**Figure 7**). α-SMA immunoreactivity was more abundant in the peri-infarct zone than in the non-infarcted and infarcted myocardium of MI hearts and the myocardium of sham hearts. However, MK2 - deficiency did not affect the distribution or intensity of α-SMA immunoreactivity. These results reveal a similar recruitment and activation of fibroblasts post-MI in hearts from wild type and MK2-deficient mice, suggesting the initiation of fibrosis was not affected by the absence of MK2.

**Figure 7.**
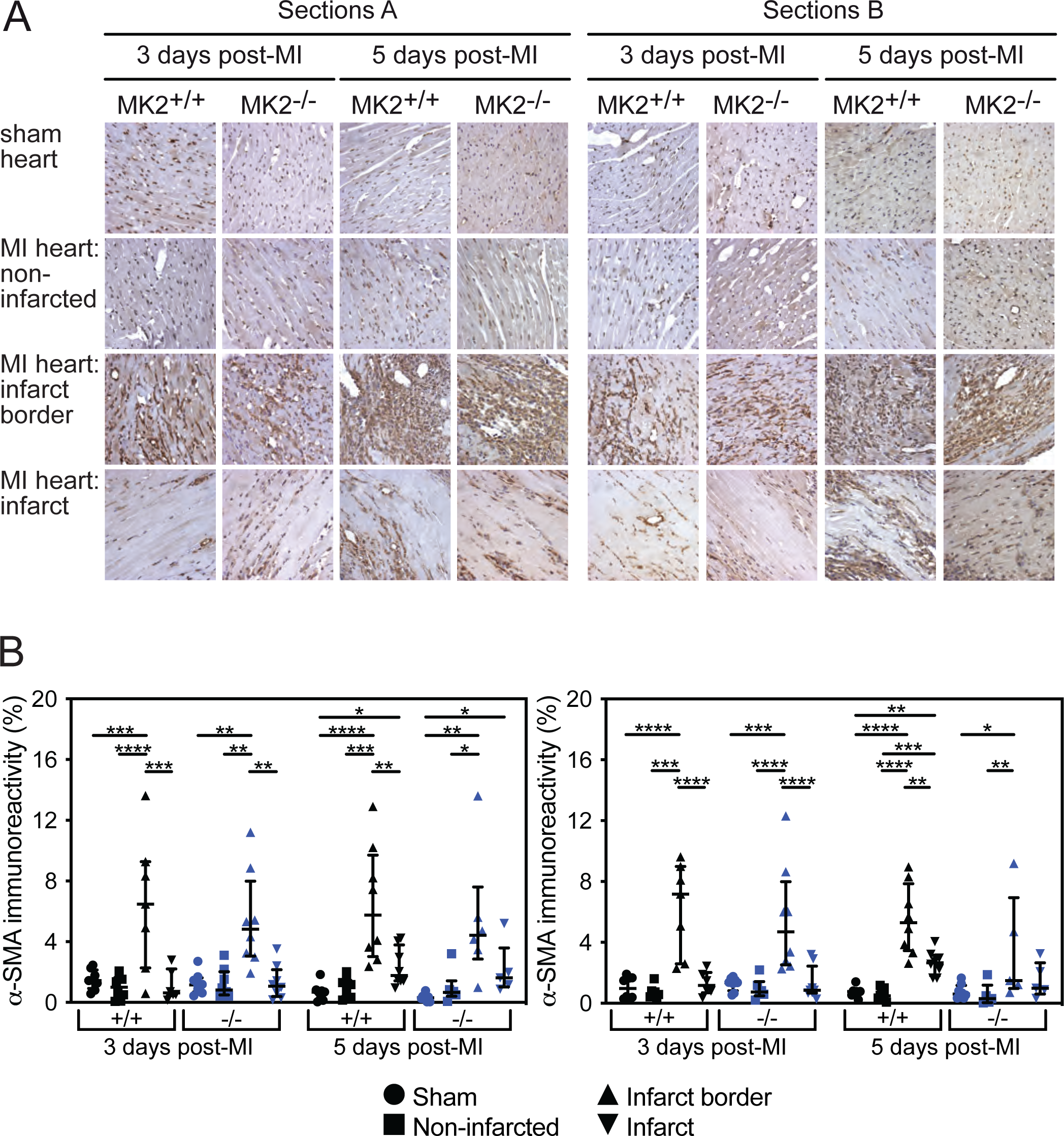
MK2-deficiency did not alter the distribution or abundance of myofibroblasts 3- and 5-days post-MI. A: Representative images of immunohistochemical staining of smooth muscle alpha-actin (α-SMA, dark brown), a myofibroblast marker, in sham and infarcted hearts from MK2^+/+^ and MK2^-/-^ mice sacrificed 3- and 5-days post-MI. Hearts were cut along the short axis through the center of the infarct to yield two pieces (Section A, upper region; Section B, lower region). **B**, α-SMA immunostaining expressed as a percentage of the total field area from sham-MK2^+/+^ (black circles, n = 7 - 8), MI-MK2^+/+^ (non-infarcted tissue: black squares, n = 7 - 8; peri-infarct: black triangles, n = 7 - 8; infarct: black inverted triangles, n = 6 - 8), sham-MK2^-/-^ (blue circles, n = 8), and MI-MK2^-/-^ (non-infarcted tissue: blue squares, n = 5 - 8; peri-infarct: blue triangles, n = 5 - 8; infarct: blue inverted triangles, n = 5 - 8) hearts. Images were analyzed by color segmentation using Image Pro Plus version 7.0 (Media Cybernetics, Silver Spring, MD). Shapiro-Wilk tests for normality were performed on all data. Data are presented as median (first quartile and third quartile). Data with a lognormal distribution were log-transformed prior to statistical comparison by two-way ANOVA, which included a factor for surgery (sham, MI), a factor for genotype (MK2^+/+^, MK2^-/-^), and a surgery x genotype interaction term. No interaction was detected (*P* > 0.05). The ANOVA was followed by Tukey’s post hoc tests for multiple comparisons. \**P* < 0.05, \*\**P* < 0.01, \*\*\**P* < 0.001, \*\*\*\**P* < 0.0001.

### Vascularization

Angiogenesis is an important component of wound repair in that it reduces myocyte mortality and maintains cardiac function by rebuilding the vasculature required to provide oxygen and nutrients to cardiac tissue ^70–72^. As deletion of MK2 impedes angiogenesis in cutaneous wound repair ^73^ and colorectal cancer ^74, 75^, we next sought to assess the effect of MK2-deficiency on angiogenesis post-MI. First, immunohistochemical staining for CD31, an endothelial cell marker ^76^, was undertaken. In general, CD31 immunoreactivity was most abundant in the peri-infarct zone in ‘A’ and ‘B’ sections for both genotypes (**Figure 8A and 8B**). On days 3 and 5 post-MI, CD31 staining was greater in the peri-infarct zone of MI-MK2^-/-^ hearts compared to that of MI-MK2^+/+^ hearts in both ‘A’ and ‘B’ sections (**Figure 8A and 8B**). In addition, 5 days post-MI, CD31 staining was greater in the infarct zone of MI-MK2^-/-^ hearts compared to that of MI-MK2^+/+^ hearts in the ‘B’ sections (**Figure 8B**) and was associated with a significant surgery x genotype interaction effect (*P* < 0.05). Thus, the absence of MK2 activity appeared to promote the recruitment of CD31-positive endothelial cells to the infarct and peri-infarct zones.

**Figure 8.**
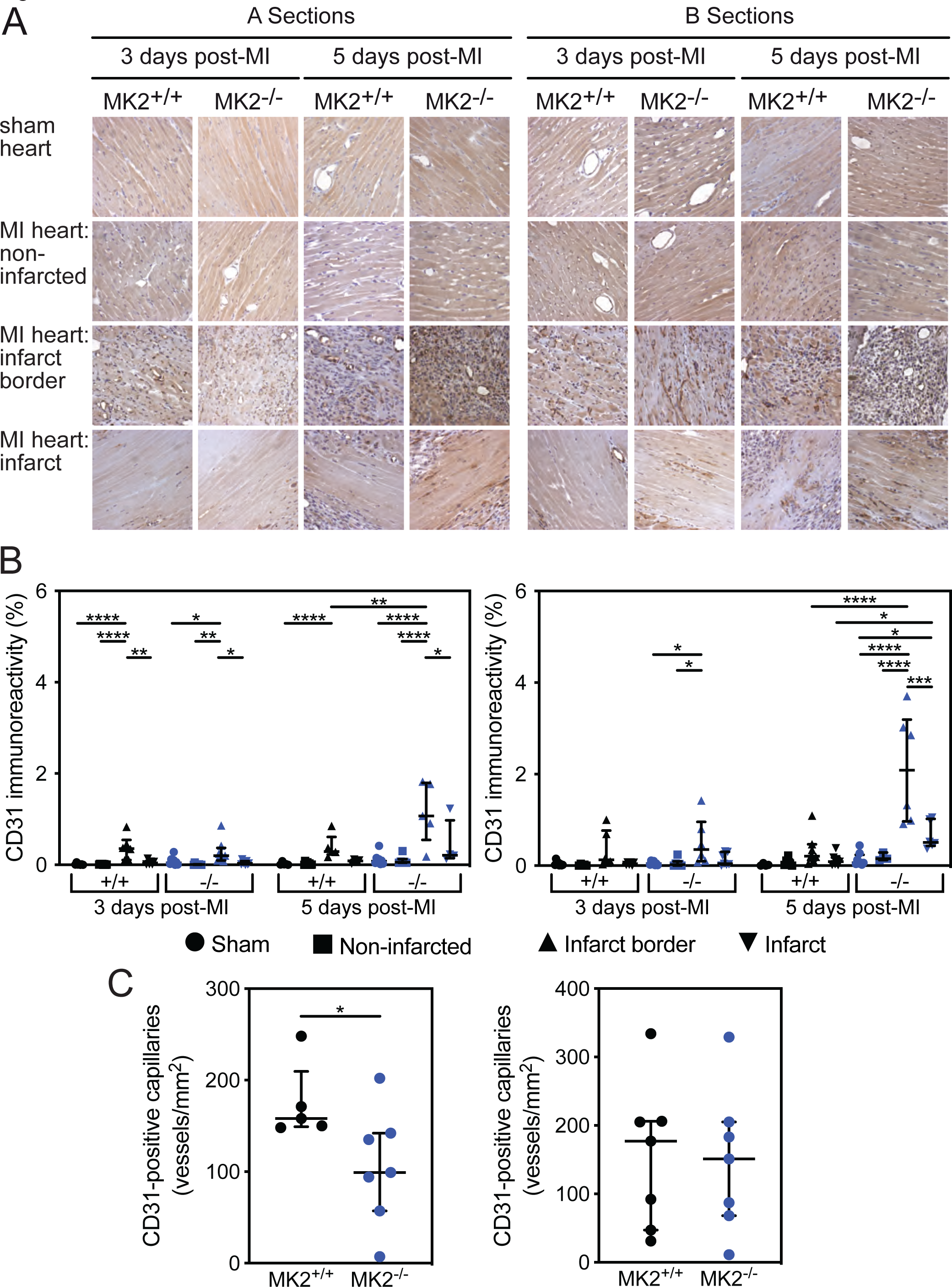
MK2-deficient hearts showed a greater increase in endothelial cell abundance in their peri-infarct region 5 days post-MI. **A**, Representative images of immunohistochemical staining of the cluster of differentiation 31 protein (CD31, dark brown), an endothelial cell marker, in MK2^+/+^ and MK2^-/-^ sham and infarct hearts collected 3- and 5-days post-MI. Hearts were cut along the short axis through the center of the infarct to yield two pieces (Section A, upper region; Section B, lower region). **B**, CD31 immunostaining expressed as a percentage of the total field area from sham-MK2^+/+^ (black circles, n = 4 - 8) and MI-MK2^+/+^ (non-infarcted tissue: black squares, n = 5 - 8; peri-infarct: black triangles, n = 5 - 7; infarct: black inverted triangles, n = 4 - 7), sham-MK2^-/-^ (blue circles, n = 4 - 8), and MI-MK2^-/-^ (non-infarcted tissue: blue squares, n = 3 - 8; peri-infarct: blue triangles, n = 5 - 8; infarct: blue inverted triangles, n = 4 - 8) hearts. Images were analyzed by color segmentation using Image Pro Plus version 7.0 (Media Cybernetics, Silver Spring, MD). Shapiro-Wilk tests for normality were performed on all data. Data are presented as median (first quartile and third quartile). Data with a lognormal distribution were log-transformed prior to statistical comparison by two-way ANOVA, which included a factor for surgery (sham, MI), a factor for genotype (MK2^+/+^, MK2^-/-^), and a surgery x genotype interaction term. A significant surgery x genotype interaction (*P* < 0.05) was detected in the ‘B’ sections 5-days post-MI. The ANOVA was followed by Tukey’s post hoc tests for multiple comparisons. \**P* < 0.05, \*\**P* < 0.01, \*\*\**P* < 0.001, \*\*\*\**P* < 0.0001. **C**, Density of new vessels as determined by the number of CD31-positive capillaries that are 20 μm in diameter or less per mm^2^ found in the peri-infarct region of MK2^+/+^ (n = 5 - 7) and MK2^-/-^ (n = 7) hearts. Data are presented as median (first quartile and third quartile). Mann-Whitney tests were performed for statistical comparisons between MK2^+/+^ and MK2^-/-^. \**P* < 0.05.

We next examined if the absence of MK2 altered the density of vessels with a diameter of 20 μm or less in the infarct border region five-days post-MI using an immunohistological approach with antibodies directed against CD31 and *α*-SMA. No *α*-SMA-positive vessels of this diameter were detected **(Data not shown)**. However, the density of CD31-positive vessels with a diameter of 20 μm or less was significantly reduced in the infarct border region of MI-MK2^-/-^ hearts, relative to MI-MK2^+/+^ hearts, in the ‘A’ sections (**Figure 8C**).

### The effects of an MK2-deficiency on cytokine production following an MI

Known substrates for MK2 include the RNA-binding proteins tristetraprolin (TTP), HuR, and ARE/poly(U)-binding/degradation factor 1 (AUF1) ^43, 49^. These proteins bind to AU-rich elements (ARE) present in the 3’-untranslated region (UTR) of numerous mRNAs, including those of several pro-inflammatory cytokines ^43, 49^, and either stabilize (HuR) or destabilize (TTP, AUF1) mRNA, implicating MK2 in the post-transcriptional regulation of gene expression ^43, 49^. MK2-mediated phosphorylation of TTP and HuR has opposing effects on their binding to mRNA: TTP dissociates from and promotes the stability of AU-rich mRNA whereas HuR binds to and stabilizes its target mRNAs ^43, 49^. Hence, we next examined the effect of MK2-deficiency on the abundance of inflammatory cytokine mRNAs in healthy and infarcted cardiac tissue 3- and 5-days post-MI using RT^2^ Profiler PCR Arrays (QIAGEN PAMM-150Z; **Supplementary Tables 1 - 16**). These data, normalized to the abundance of each transcript in the ventricular myocardium of wild type sham mice, are summarized in the form of volcano plots in **Figure 9**. Due to space limitations, some, but not all, mRNAs that underwent a 2-fold change in abundance with *P* < 0.05 are identified in the figure. The inflammatory phase of repair starts within 12 hours of MI and can last up to 6 days ^3, 4, 6^. In mice, day 3 post-surgery corresponds roughly to the midpoint of the inflammatory phase and the start of granulation tissue formation whereas day 5 corresponds to the resolution of the inflammatory phase. Both the absence of MK2 and time after MI affected the abundance of numerous transcripts in the sham, infarcted, and non-infarcted myocardium. It is also worth noting that the abundance of *Il15* mRNA was reduced in MK2-deficient sham hearts (**Figure 9**).

**Figure 9.**
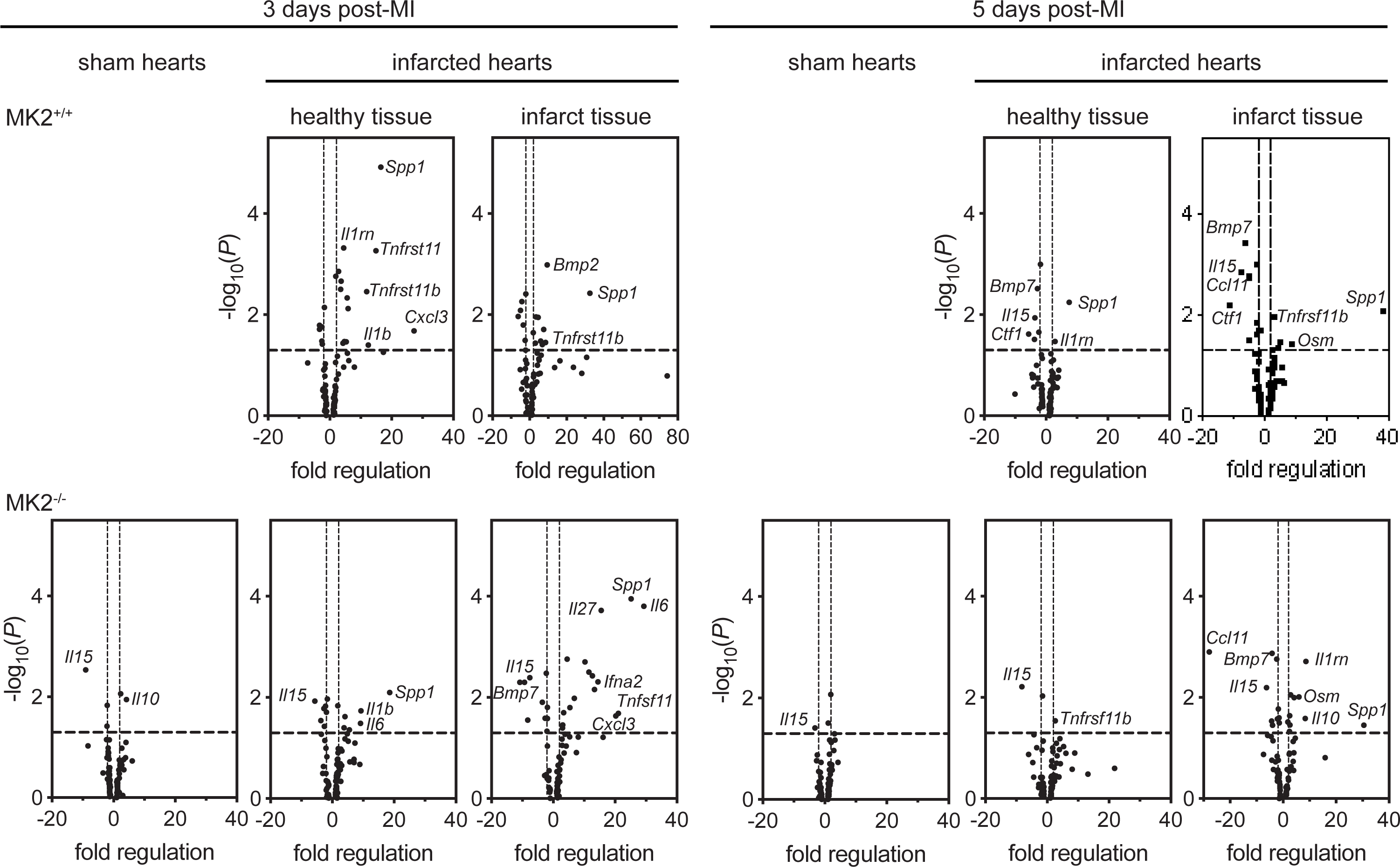
MK2-deficiency alters transcript expression pattern but does not impair overall inflammatory response. Volcano plots showing the abundance of inflammatory transcripts in sham, healthy and infarct left ventricular (LV) tissue from MK2^+/+^ and MK2^-/-^ mice 3- and 5-days post-MI (n = 3 to 4). mRNA abundance was quantified using RT^2^ Profiler PCR Arrays (QIAGEN PAMM-150Z). Each transcript was normalized to its abundance in sham-MK2^+/+^ LV tissues (n = 3 - 4) at day 3 or day 5 post-MI. Some transcripts that underwent significant changes have been labeled. Transcripts above the horizontal dotted line showed a fold change with *P* < 0.05. Transcripts found outside of the vertical dotted lines underwent a 2-fold change in abundance (left, decreased abundance; right, increased abundance).

The absence of MK2 impairs the induction of several cytokines in cells and mice in response to lipopolysaccharide (LPS) ^46^. A comparison of the cytokine transcripts that have been shown to be increased in response to an LPS challenge and attenuated by the absence of MK2 shows that many of those were not elevated 3- **(Supplementary Table 15)** or 5-days (**Supplementary Table 16**) post-MI. However, an important caveat to this comparison is that the time courses of the LPS studies are much shorter than 3-5 days. To examine the effects of MK2-deficiency more closely, the array data from MI-MK2^-/-^ mice was normalized to the corresponding dataset from MI-MK2^+/+^ mice (**Table 3**) and replotted as volcano plots (**Figure 10**). Focusing on mRNAs that underwent a 2-fold change in abundance with *P* < 0.05, Figure 10 shows that at both 3- and 5-days post-MI, the absence of MK2 reduced the abundance of some mRNAs but increased the abundance of others. Furthermore, the effects of MK2-deficiency on transcript abundance differed both with respect to surviving versus infarcted myocardium and time post-MI. Three days post-MI, *Ifna2* was increased and *Il16* was decreased in infarcted MK2^-/-^ tissue whereas in the non-infarcted myocardium *Il27* increased and *Tnfsf11*, *Ccl3*, and *Il1rn* decreased. Five days post-MI, *Ctf16* and *Il10* were increased in infarcted tissue whereas in the non-infarcted myocardium *Ccl9, Nodal, and Xcl2* increased and *Il15* decreased. Furthermore, as shown in Figure 9, in sham-operated mice 3 days post-surgery, *Il15* was reduced and *Il10* was increased in MK2-deficient hearts. Five-days post-surgery, *Il15* was decreased. These findings show that the absence of MK2 did modify the inflammatory response in the infarcted heart by both potentiating and attenuating the induction of various cytokines.

**Table 3.**
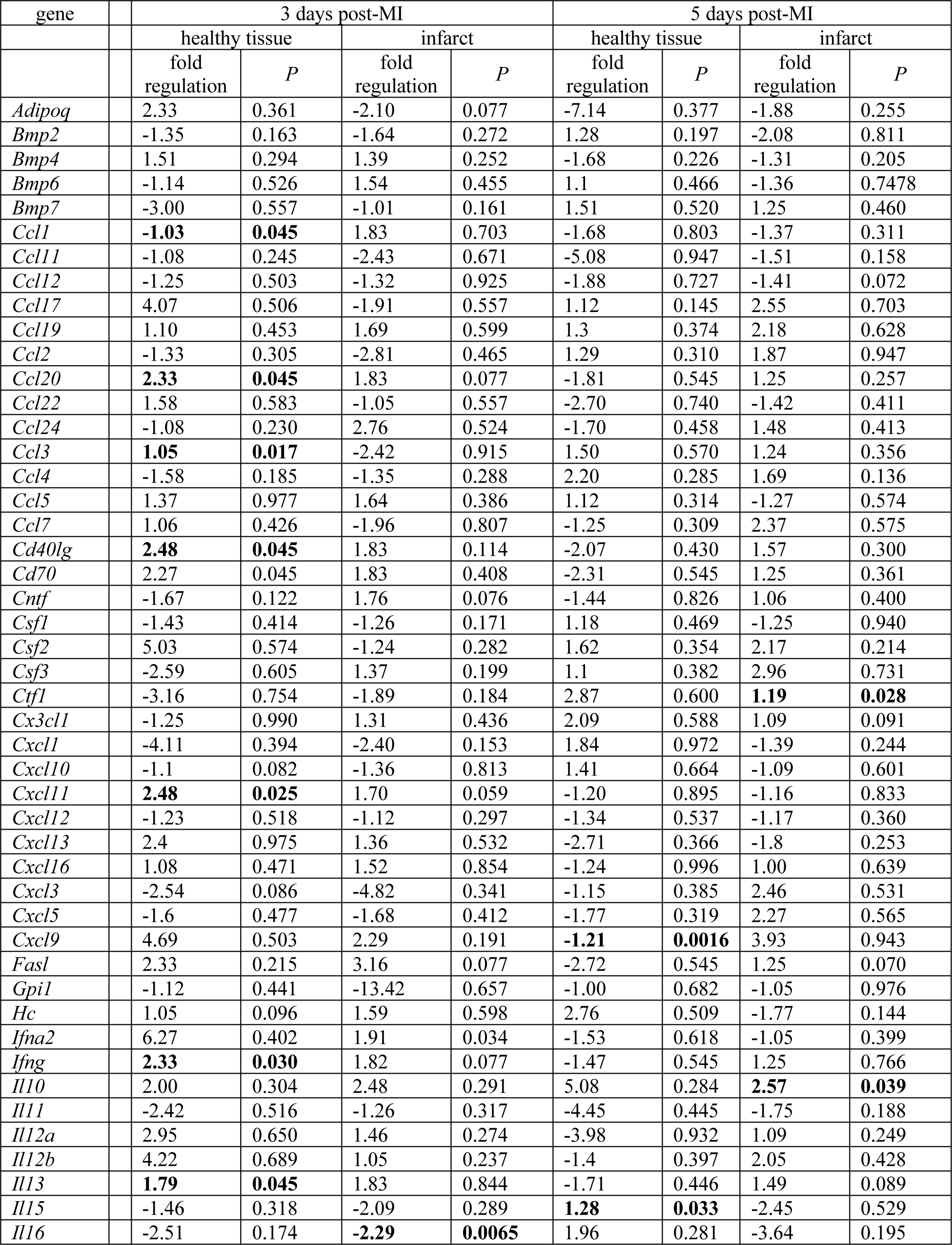
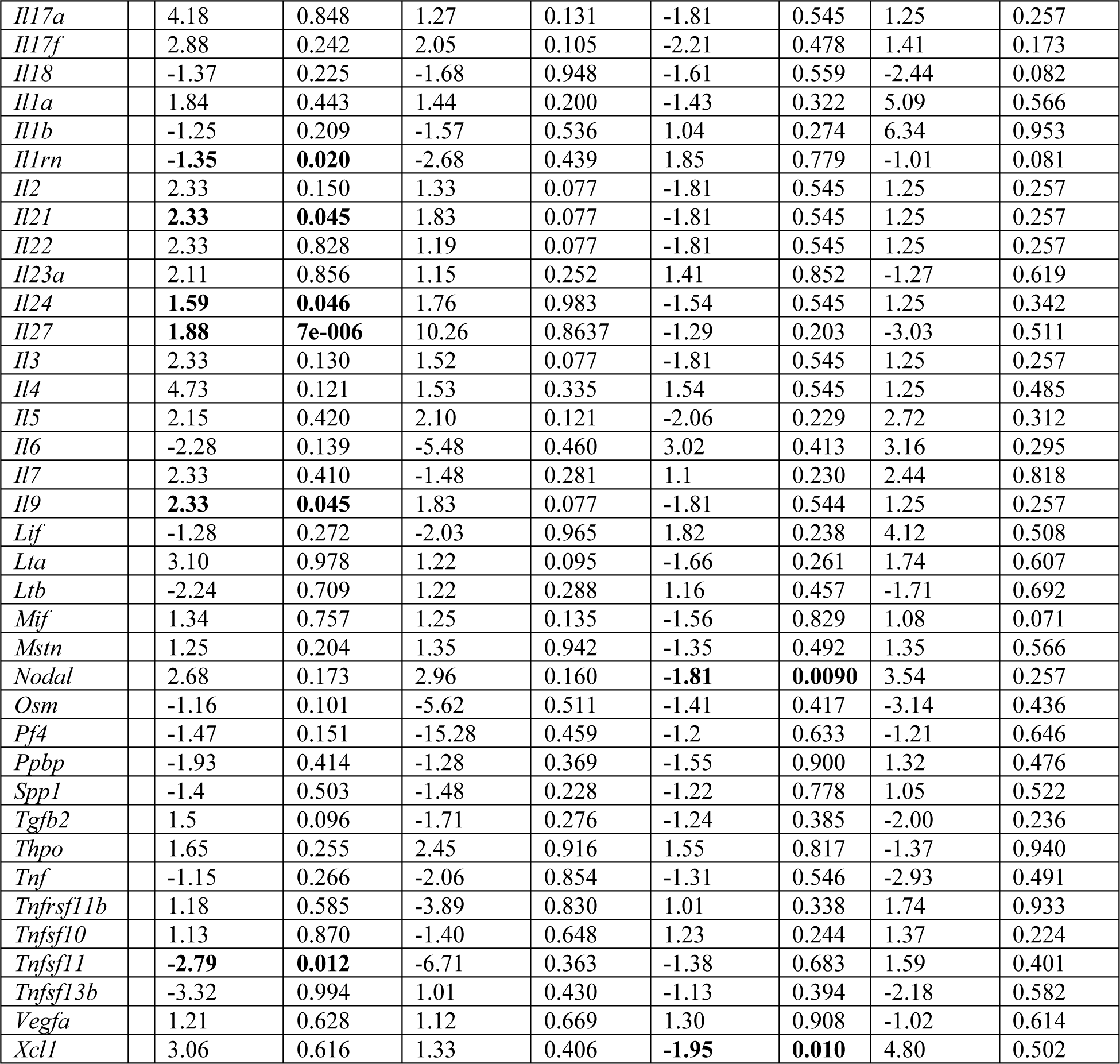
Effect of MK2-deficiency on cytokine transcript abundance following MI. Data shown are expressed as the fold-regulation in transcript abundance relative to the respective values from MK2^+/+^ hearts. Fold-regulation: Fold-change values greater than one indicate an increase in transcript abundance, relative to that of MK2^+/+^ cardiac tissue, and the fold-regulation is equal to the fold-change. Where the transcript abundance is less than that of MK2^+/+^ cardiac tissue, the fold-change is less than one and the fold-regulation is the negative inverse of the fold-change. Cells where P < 0.05 are indicated in bold. n = 3 - 4.

**Figure 10.**
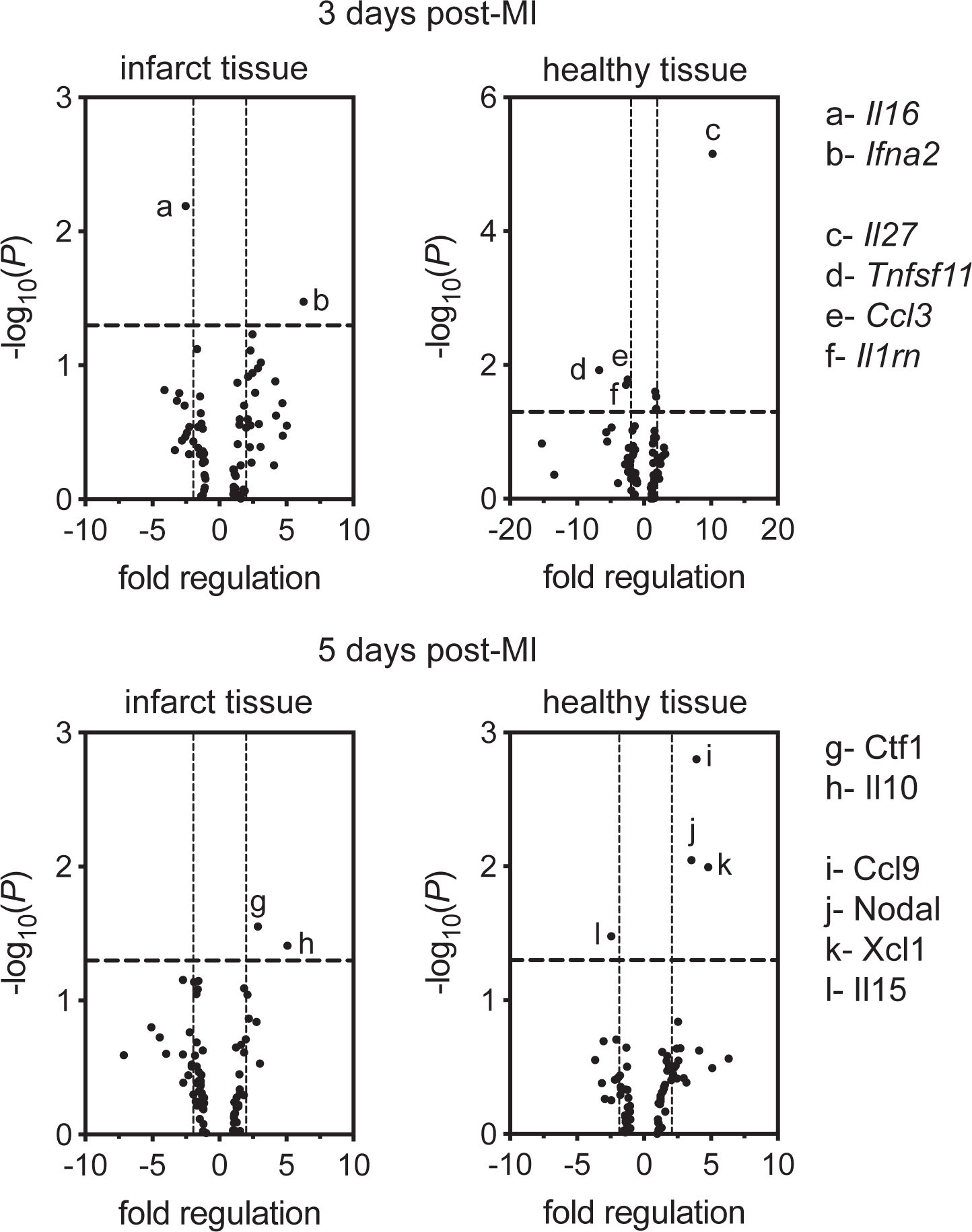
MK2-deficiency leads to a different pro- and anti-inflammatory signature in both infract and healthy tissues at 3- and 5-days post-MI. Volcano plots representing the abundance of inflammatory transcripts in MK2-deficient infarct or healthy tissues normalized to the corresponding MK2^+/+^ heart tissue (n = 3 to 4). mRNA abundance was quantified using RT^2^ Profiler PCR Arrays (QIAGEN PAMM-150Z). Transcripts that underwent a significant change are labeled. Transcripts above the horizontal dotted line showed a fold change with *P* < 0.05. Transcripts found outside of the vertical dotted lines underwent a 2-fold change in abundance (left, decreased abundance; right, increased abundance).

### MK2-deficiency did not alter myofibroblast motility in vitro

HSP27/25, a small heat shock protein with actin-capping activity, is a substrate for MK2 ^36, 44^ and MK2-deficiency in immortalized MEFs, tracheal smooth muscle cells, endothelial cells, and macrophages decreases migration ^77–79^. Since fibroblast recruitment is an important component of wound healing post-MI, we next examined migration by scratch-wound assay in ventricular fibroblasts isolated from the hearts of MK2^+/+^ and MK2^-/-^ mice. Fibroblasts were grown to 80% confluence on a rigid plastic substrate, “wounds” were created, and the cells were incubated for an additional 24 h with or without the addition of serum alone or serum plus angiotensin II to the media. Wound areas were measured morphometrically at time 0 and after 24 h. The percentage of wound closure was comparable between genotypes under all 3 experimental conditions (**Figure 11**). Hence, myofibroblast migration *in vitro* was not altered by the absence of MK2.

**Figure 11.**
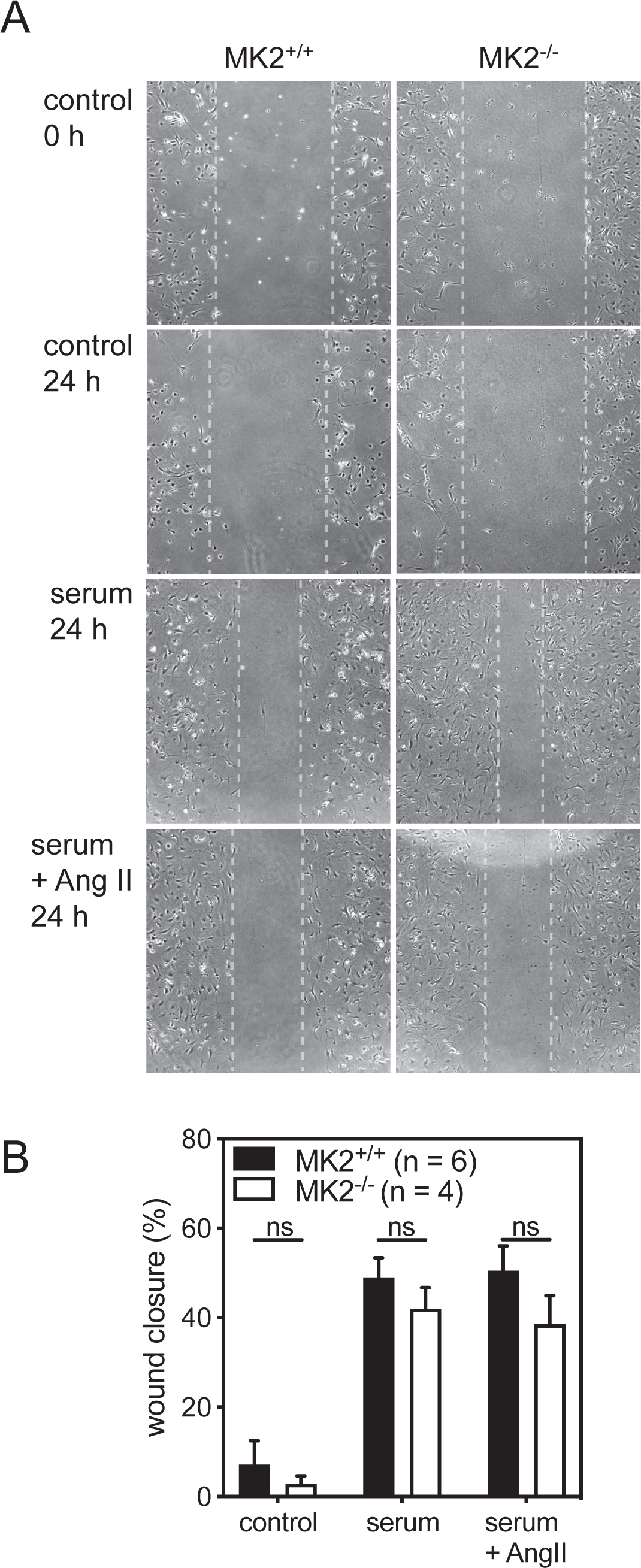
MK2-deficiency does not alter myofibroblast motility *in vitro*. **A**, Representative images of scratch wound assays of left ventricular myocardial myofibroblasts isolated from MK2^+/+^ and MK2^-/-^ mice. Scratches were imposed once cell confluency reached 80%. Cells were then incubated for 24 h with serum-free media (control), media supplemented with 10% serum, or media supplemented with 10% serum and angiotensin II (Ang II, 1 μM). Open wound area (space between vertical dotted lines) was measured at times 0 and 24 h and wound closure was calculated. **B**, Motility, expressed as percentage of wound area remaining open after 24 h. Data was normalized to the values for MK2^+/+^ myofibroblasts maintained in serum-free media (6.26 ± 16.5%) and are presented as mean ± SEM (n = 4). MK2^+/+^, solid bars; MK2^+/+^, open bars. Statistical analysis was done by 2-way ANOVA followed by Tukey’s post hoc test.

### Discussion

MK2 plays a prominent role in the inflammatory response via both regulating the stability of many cytokine mRNAs and mediating the response of numerous cytokine receptors ^80^. As inflammation is an important early phase of myocardial wound healing post-myocardial infarct, we examined the effects of MK2-deficiency in 3-month-old male mice 3- and 5-days after permanent ligation of the left anterior descending coronary artery (LAD). Mortality was reduced in MK2-deficient mice as the incidence of left ventricular (LV) wall rupture during the first 5 days of wound-healing post-MI was lower in MI-MK2^-/-^ than MI-MK2^+/+^ mice. However, MK2-deficiency did not affect infarct area or immune cell infiltration whereas LV dilation was attenuated in MI-MK2^-/-^ mice. Hence, the MK2-deficiency did not impair the inflammatory phase of post-MI wound repair.

Wound healing post-MI progresses through three overlapping phases: inflammation, proliferation, and maturation ^3, 5^ and involves components of the innate immune system. Within the infarct, necrotic myocytes release damage-associated molecular patterns (DAMPs), resulting in the induction and release of pro-inflammatory cytokines and chemokines and the subsequent recruitment of neutrophils and monocytes. Monocytes differentiate into macrophages and, together, these cells phagocytose dead tissue and release inflammatory mediators. After several days, the inflammatory response begins to resolve. Appropriate induction and resolution of the inflammatory response is an important step prior to fibroblast activation, secretion of extracellular matrix proteins, and formation of a mature scar. MK2-deficiency can affect multiple aspects of the inflammatory response, including immune cell recruitment ^47, 55, 81–83^, macrophage activation ^47, 84, 85^, macrophage M1/M2 polarization ^52, 74^, cell motility ^77, 81^, cytokine/chemokine production ^46, 47, 77, 81, 83, 85, 86^, and signaling downstream of cytokine receptors ^87–89^. The present study employed a global knockout model ^77^ to study the effects of an MK2-deficiency on the inflammatory phase of wound repair post-MI. This study examined mice on days 3 and 5 post-MI, as the primary cause of death in mice following LAD ligation is rupture of the LV wall at the infarct border on or around day 4 ^63^. The area at risk and infarct area in MI-MK2^-/-^ and MI-MK2^+/+^ mice were similar. However, whereas fibroblast-targeted deletion of p38*α*, which, along with p38*β*, serves to activate MK2, resulted in 100% mortality following permanent LAD ligation ^26^, survival was actually increased in MK2-deficient mice five-days post-MI. Hence, the absence of MK2 activity was not detrimental overall to the inflammatory phase of wound-repair post-MI.

In response to pro-inflammatory mediators released by necrotic and apoptotic myocytes in the infarcted myocardium, circulating neutrophils and monocytes migrate to the infarct ^20, 30^ where they, along with resident fibroblasts, degrade the extracellular matrix, remove dead cells and debris, and participate in the inflammatory response by secreting numerous cytokines ^27, 64, 68^. MK2 inhibition or deficiency has been shown to reduce myeloid cell recruitment ^47, 55, 81–83^, which likely results from the roles played by MK2 in cytokine/chemokine production, the cellular signaling downstream of cytokine/chemokine receptor activation, and cell motility. However, in the context of MI, MK2-deficiency did not alter the intensity of myeloperoxidase (MPO) immunostaining in the infarct or infarct border region, suggesting recruitment of neutrophils or monocytes was unaffected 3- or 5-days post-MI. This study would have missed changes in neutrophil recruitment, which peaks 1-day post-MI, and would not discriminate between Ly-6C^hi^ and Ly-6C^lo^ monocytes, which peak on day-3 and day-7 post-MI, respectively ^30^. Ly-6C^hi^ monocytes are pro-inflammatory whereas Ly-6C^lo^ monocytes have been implicated in resolving the inflammatory response ^30, 90–92^. In an inflammatory environment such as the infarct zone, monocytes differentiate into macrophages, supplementing the complement of resident macrophages ^93, 94^. In the healthy myocardium, resident macrophages comprise both pro-inflammatory/M1 and anti-inflammatory/M2 phenotypes ^94^ where they phagocytose debris and apoptotic cells. Initially following MI, M1 macrophages predominate, contributing to acute inflammation and phagocytosis. As the inflammatory response progresses macrophages polarize towards the M2 phenotype, which are implicated in resolution of the inflammatory response and promote fibrosis. This polarization depends, in part, on the secretion of neutrophil gelatinase-associated lipocalin (NGAL) ^95^. An MK2-deficiency does not affect the abundance of circulating CD14^+^ monocytes/macrophages ^46^, whereas macrophage recruitment and activation are reduced ^47, 77, 82, 85, 96^. In addition, in a mouse model of inflammation-driven colorectal cancer ^75^ and compression-induced spinal cord injury ^97^, an MK2-deficiency reduces M2 polarization. Similarly, IL4/IL-13-induced M2 polarization of human U-837 monocytic cell-derived macrophages is reduced by the MK2 inhibitor PF 3644022 ^75^. However, the abundance of M2 macrophages was not significantly reduced in the infarct or peri-infarct regions of MI-MK2^-/-^ hearts post-MI. Although administration of MMI-0100, a cell-permeable peptide inhibitor of MK2, was previously shown to improve systolic function post-MI ^98, 99^, in the present study, systolic function in MI-MK2^-/-^ mice did not differ significantly from MI-MK2^+/+^ mice. Furthermore, dilation was actually reduced in MI-MK2^-/-^ hearts, which suggests the innate immune response was unimpaired, and an excessive inflammatory response had not occurred.

The secretion of pro-inflammatory cytokines within the infarcted myocardium results in fibroblast recruitment, proliferation, and activation to a myofibroblast phenotype (see ^100^). Myofibroblasts then secrete extracellular matrix proteins to create a scar and reinforce the damaged ventricular wall. Although an MK2-deficiency impairs motility in mouse embryonic fibroblasts ^77^, the motility of adult mouse ventricular myofibroblasts was not affected by the absence of MK2. In addition, the abundance of myofibroblasts in the infarct or peri-infarct zone was not affected by MK2-deficiency and, although early in the reparative process, only small differences were observed in the collagen content within the infarct. Taken together, these observations suggest that fibroblast activation was not affected by the absence of MK2 at least up until day 5 post-MI. The effects of MK2-deficiency on the proliferative and maturation phases of wound repair post-MI remain to be determined.

The coordinated production of pro- and anti-inflammatory cytokines orchestrates the onset and resolution of the inflammatory phase of cardiac repair post-MI. Changes in cytokine mRNA, assessed using pathway-targeted qPCR arrays (QIAGEN) and normalized to their respective sham wild type levels, indicated that infarction resulted in an inflammatory response in both MK2^+/+^ and MK2^-/-^ hearts. Indeed, the increased abundance of *Tgfb*2 mRNA, an anti-inflammatory cytokine, as well as the decreased abundance of transcripts for *Bmp7*, a *Tgfb2* antagonist, and *Il15*, a pro-inflammatory cytokine, were observed in infarcted MK2^+/+^ and MK2^-/-^ tissue 3 days post-MI ^101–107^. In addition, the abundance of *Il11* mRNA, a cytokine secreted by fibroblasts in response to TGF-*β*, was also increased in infarcts of both genotypes ^108, 109^. A decreased abundance of *Il15* mRNA is also detected in all tissues 5 days post-MI. Previous studies observed that the absence of MK2 reduces the abundance of mRNA for numerous cytokines when an inflammatory response is evoked ^46^. Normalizing qPCR array data from MK2-deficient tissue to the corresponding wild type tissue revealed that, 3-days post-MI, the abundance of *Ifna2* mRNA was increased and *Il16* was decreased in infarcted tissue whereas in the non-infarcted myocardium *Il27* increased and *Tnfsf11*, *Ccl3*, and *Il1rn* were decreased in MK2-deficient tissue relative to wild type. Five days post-MI, the abundance of *Ctf16* and *Il10* increased in MK2-deficient infarcted tissue whereas in the non-infarcted myocardium *Ccl9, Nodal, and Xcl2* increased and *Il15* decreased. Hence, during the inflammatory phase of wound repair following MI, the absence of MK2 resulted in both increases and decreases in cytokine mRNA.

### Conclusions

In conclusion, the present study shows that, although the mRNA levels for several cytokines were affected, an MK2-deficiency did not impair the inflammatory phase of wound healing following myocardial infarction. In fact, rather than increased mortality, mice deficient in MK2 showed reduced dilation and a significantly higher survival rate than wild type mice. It remains to be determined if a deficiency in MK2 is detrimental to collagen deposition or scar maturation.

## Additional information

### Author contributions

JT, SAN, PS, MGS, and BGA conceived and designed the experiments. JT, SAN, PS, FS, ND, DG, MAG, YS, CT, and MECL performed the experiments and analyzed the data. JT and BGA assembled and interpreted all the data. MG provided the mouse model. JT and BGA wrote and critically reviewed the manuscript. All authors have approved the final version of the current manuscript.

## Acknowledgements

We thank Ms. Karine Nadeau for animal care and breeding.

