## Supplementary Table 1 for "MK2 deficiency decreases mortality during the inflammatory phase after myocardial infarction in mice"

**Supplementary Table 1. RT2 profiler PCR array analysis of anti-inflammatory cytokine mRNA in mouse left ventricular tissue 3 days post-MI.**

| Anti-inflammatory cytokine | | MK2^+/+^ | | MK2^-/-^ | | |
| --- | --- | --- | --- | --- | --- | --- |
| Symbol | Official full name | Infarct tissues | Healthy tissues | Sham | Infarct tissues | Healthy tissues |
| ***Ccl19*** | **chemokine (C-C motif) ligand 19** | **-2.34 (0.032)** | -1.49 (0.318) | -1.59 (0.129) | **-2.03 (0.047)** | 1.34 (0.353) |
| *Il2* | interleukin 2 | -1.35 (0.388) | 1.30 (0.576) | 1.08 (0.793) | 1.18 (0.724) | 1.38 (0.350) |
| *Il4* | interleukin 4 | -2.27 (0.215) | -1.53 (0.356) | -1.65 (0.339) | 1.37 (0.573) | -1.22 (0.831) |
| ***Il6*** | **interleukin 6** | 23.7 (0.110) | 17.4 (0.055) | 1.19 (0.964) | **29.3 (0.0002)** | **9.16 (0.033)** |
| ***Il10*** | **interleukin 10** | 4.69 (0.149) | 5.71 (0.058) | **4.16 (0.011)** | 7.49 (0.123) | 8.88 (0.211) |
| ***Il11*** | **interleukin 11** | 5.64 (0.145) | 1.96 (0.277) | -1.37 (0.421) | **3.03 (0.035)** | 1.36 (0.383) |
| *Il12a* | interleukin 12a | -5.02 (0.122) | -7.15 (0.090) | -8.29 (0.094) | -1.47 (0.599) | -2.05 (0.317) |
| *Il12b* | interleukin 12b | -1.35 (0.388) | 1.85 (0.329) | 2.67 (0.225) | 4.22 (0.092) | 1.95 (0.103) |
| *Il13* | interleukin 13 | -1.23 (0.843) | -1.61 (0.413) | -1.21 (0.679) | -1.08 (0.640) | 1.53 (0.946) |
| ***Il18*** | **interleukin 18** | 1.76 (0.446) | **5.93 (0.008)** | 1.35 (0.688) | 2.00 (0.357) | **3.82 (0.040)** |
| *Il22* | interleukin 22 | -1.35 (0.388) | 1.57 (0.383) | -1.39 (0.295) | 1.13 (0.778) | 1.38 (0.350) |
| *Il23a* | interleukin 23, alpha subunit p19 | -2.13 (0.202) | 1.25 0.598 | -1.43 (0.535) | -1.01 (0.985) | -1.17 (0.549) |
| *Il24* | interleukin 24 | 1.28 (0.447) | -1.07 (0.616) | -1.39 (0.295) | 1.33 (0.481) | 1.39 (0.348) |
| ***Tgfb2*** | **transforming growth factor, beta 2** | **2.04 (0.023)** | **1.91 (0.002)** | -1.43 (0.071) | **3.01 (0.040)** | 1.01 (0.827) |

Data shown are expressed as the fold-regulation in transcript abundance relative to LV tissue from sham MK2^+/+^ mice. Fold-regulation: Fold-change values greater than one indicate an increase in transcript abundance, relative to that of LV tissue from sham MK2^+/+^ mice, and the fold-regulation is equal to the fold-change. Where the transcript abundance is less than that of LV tissue from sham MK2^+/+^ mice, the fold-change is less than one and the fold-regulation is the negative inverse of the fold-change. *P*-values are indicated in parentheses. *N* = 3 or 4 (MK2^+/+^ sham).
