## Supplementary Table 2 for "MK2 deficiency decreases mortality during the inflammatory phase after myocardial infarction in mice"

**Supplementary Table 2. RT2 profiler PCR array analysis of anti-inflammatory cytokine mRNA in mouse left ventricular tissue 5 days post-MI.**

| Anti-inflammatory cytokine | | MK2^+/+^ | | MK2^-/-^ | | |
| --- | --- | --- | --- | --- | --- | --- |
| Symbol | Official full name | Infarct tissues | Healthy tissues | Sham | Infarct tissues | Healthy tissues |
| *Ccl19* | chemokine (C-C motif) ligand 19 | -1.38 (0.723) | -1.22 (0.468) | 1.60 (0.198) | -1.16 (0.625) | 1.91 (0.396) |
| *Il2* | interleukin 2 | 1.38 (0.818) | 1.21 (0.906) | 1.07 (0.980) | -1.35 (0.465) | 1.62 (0.616) |
| *Il4* | interleukin 4 | 1.56 (0.605) | 1.21 (0.906) | 1.70 (0.478) | 2.30 (0.288) | 1.62 (0.616) |
| *Il6* | interleukin 6 | 5.64 (0.113) | 4.00 (0.173) | 1.35 (0.513) | 15.9 (0.156) | 13.2 (0.328) |
| ***Il10*** | **interleukin 10** | 1.78 (0.342) | 2.26 (0.225) | 3.27 (0.068) | **8.38 (0.026)** | 6.07 (0.127) |
| *Il11* | interleukin 11 | 5.38 (0.203) | 3.59 (0.279) | 1.32 (0.628) | 1.18 (0.746) | 2.24 (0.243) |
| *Il12a* | interleukin 12a | 1.89 (0.302) | 1.48 (0.530) | 1.76 (0.318) | -2.12 (0.449) | 1.71 (0.377) |
| *Il12b* | interleukin 12b | -1.36 (0.373) | -1.41 (0.360) | 1.31 (0.540) | -1.98 (0.271) | 1.56 (0.498) |
| *Il13* | interleukin 13 | 1.16 (0.747) | 1.48 (0.730) | 1.64 (0.480) | -1.53 (0.387) | 2.41 (0.405) |
| *Il18* | interleukin 18 | 1.95 (0.224) | 1.14 (0.990) | -1.17 (0.952) | 1.16 (0.862) | -1.96 (0.828) |
| *Il22* | interleukin 22 | 1.38 (0.818) | 1.21 (0.906) | 1.43 (0.690) | -1.35 (0.465) | 1.62 (0.616) |
| *Il23a* | interleukin 23, alpha subunit p19 | -2.61 (0.025) | -1.45 (0.428) | -1.01 (0.801) | -1.89 (0.353) | -1.77 (0.750) |
| *Il24* | interleukin 24 | 1.38 (0.818) | 1.21 (0.906) | 1.06 (0.991) | -1.17 (0.580) | 1.62 (0.616) |
| *Tgfb2* | transforming growth factor, beta 2 | 1.16 (0.747) | 1.48 (0.730) | 1.64 (0.480) | -1.53 (0.387) | 2.41 (0.405) |

Data shown are expressed as the fold-regulation in transcript abundance relative to LV tissue from sham MK2^+/+^ mice. Fold-regulation: Fold-change values greater than one indicate an increase in transcript abundance, relative to that of LV tissue from sham MK2^+/+^ mice, and the fold-regulation is equal to the fold-change. Where the transcript abundance is less than that of LV tissue from sham MK2^+/+^ mice, the fold-change is less than one and the fold-regulation is the negative inverse of the fold-change. *P*-values are indicated in parentheses. *N* = 3 or 4 (MK2^+/+^ infarct tissue).
