## Supplementary Table 3 for "MK2 deficiency decreases mortality during the inflammatory phase after myocardial infarction in mice"

**Supplementary Table 3. RT2 profiler PCR array analysis of chemokine mRNA in mouse left ventricular tissue 3 days post-MI.**

| Chemokine | | MK2^+/+^ | | MK2^-/-^ | | |
| --- | --- | --- | --- | --- | --- | --- |
| Symbol | Official full name | Infarct tissues | Healthy tissues | Sham | Infarct tissues | Healthy tissues |
| *Ccl1* | chemokine (C-C motif) ligand 1 | 1.65 (0.272) | -1.11 (0.562) | -1.39 (0.295) | 1.13 (0.778) | 1.38 (0.350) |
| ***Ccl2*** | **chemokine (C-C motif) ligand 2** | **7.59 (0.020)** | **4.72 (0.034)** | -1.30 (0.605) | **5.35 (0.016)** | 2.63 (0.204) |
| ***Ccl3*** | **chemokine (C-C motif) ligand 3** | 5.12 (0.079) | **3.67 (0.002)** | 1.07 (0.927) | 3.83 (0.053) | 1.50 (0.212) |
| ***Ccl4*** | **chemokine (C-C motif) ligand 4** | **4.61 (0.011)** | **2.81 (0.001)** | -1.44 (0.524) | 2.84 (0.067) | 1.73 (0.175) |
| *Ccl5* | chemokine (C-C motif) ligand 5 | -1.02 (0.902) | 1.39 (0.542) | -1.30 (0.517) | 1.40 (0.541) | 1.03 (0.914) |
| ***Ccl7*** | **chemokine (C-C motif) ligand 7** | **6.94 (0.039)** | **5.57 (0.005)** | -1.20 (0.833) | 8.10 (0.061) | 4.14 (0.066) |
| *Cl11* | chemokine (C-C motif) ligand 11 | 1.28 (0.605) | 1.69 (0.256) | 1.20 (0.638) | 1.50 (0.407) | -1.77 (0.634) |
| ***Ccl12*** | **chemokine (C-C motif) ligand 12** | 5.02 (0.075) | **5.32 (0.034)** | 1.73 (0.282) | **6.81 (0.010)** | 7.35 (0.080) |
| *Ccl17* | chemokine (C-C motif) ligand 17 | -1.91 (0.884) | 1.26 (0.647) | -1.13 (0.555) | 1.59 (0.890) | -1.27 (0.573) |
| ***Ccl19*** | **chemokine (C-C motif) ligand 19** | **-2.34 (0.032)** | -1.49 (0.318) | -1.59 (0.129) | **-2.03 (0.047)** | 1.34 (0.353) |
| *Ccl20* | chemokine (C-C motif) ligand 20 | -1.35 (0.388) | -1.11 (0.562) | -1.39 (0.295) | 1.85 (0.151) | 1.38 (0.350) |
| *Ccl22* | chemokine (C-C motif) ligand 22 | 1.49 (0.748) | 1.10 (0.750) | 2.95 (0.906) | 1.56 (0.688) | 1.25 (0.560) |
| *Ccl24* | chemokine (C-C motif) ligand 22 | 1.60 (0.381) | -1.31 (0.427) | 1.04 (0.994) | 2.03 (0.273) | 3.07 (0.222) |
| *Cx3cl1* | chemokine (C-X3-C motif) ligand 1 | 1.31 (0.334) | -1.19 (0.981) | -1.23 (0.957) | -1.08 (0.646) | -1.17 (0.530) |
| *Cxcl1* | chemokine (C-X-C motif) ligand 1 | 6.31 (0.064) | 1.79 (0.493) | 1.85 (0.411) | 2.06 (0.465) | 1.53 (0.972) |
| ***Cxcl3*** | **chemokine (C-X-C motif) ligand 3** | 74.3 (0.163) | **27.3 (0.021)** | 2.55 (0.194) | **20.2 (0.023)** | **5.57 (0.044)** |
| ***Cxcl5*** | **chemokine (C-X-C motif) ligand 5** | 16.4 (0.082) | 7.92 (0.109) | 3.32 (0.278) | **13.4 (0.007)** | 5.66 (0.188) |
| *Cxcl9* | chemokine (C-X-C motif) ligand 9 | -1.35 (0.388) | -1.11 (0.562) | 1.82 (0.336) | 3.49 (0.125) | 2.62 (0.269) |
| ***Cxcl10*** | **chemokine (C-X-C motif) ligand 10** | **3.55 (0.011)** | **3.48 (0.003)** | 1.04 (0.962) | 2.88 (0.089) | 2.18 (0.112) |
| *Cxcl11* | chemokine (C-X-C motif) ligand 11 | -1.35 (0.388) | 1.17 (0.794) | 2.06 (0.229) | 1.29 (0.564) | 1.81 (0.165) |
| ***Cxcl12*** | **chemokine (C-X-C motif) ligand 12** | -1.53 (0.093) | -1.44 (0.094) | **-2.08 (0.015)** | **-1.90 (0.016)** | **-1.85 (0.020)** |
| *Cxcl13* | chemokine (C-X-C motif) ligand 13 | -1.97 (0.502) | 1.21 (0.619) | 2.12 (0.240) | 1.49 (0.723) | 1.91 (0.333) |
| ***Cxcl16*** | **chemokine (C-X-C motif) ligand 16** | **3.11 (0.037)** | 2.44 (0.066) | 1.27 (0.600) | **3.36 (0.020)** | 2.77 (0.105) |
| *Pf4* | platelet factor 4 | 2.21 (0.149) | 1.88 (0.088) | -1.76 (0.123) | 1.28 (0.482) | 1.09 (0.895) |
| *Ppbp* | pro-platelet basic protein | 2.92 (0.241) | 1.60 (0.537) | -1.72 (0.313) | 1.66 (0.495) | 1.03 (0.764) |
| *Xcl1* | chemokine (C motif) ligand 1 | -1.83 (0.531) | -2.02 (0.418) | -3.42 (0.325) | -1.98 (0.400) | -1.98 (0.439) |

Data shown are expressed as the fold-regulation in transcript abundance relative to LV tissue from sham MK2^+/+^ mice. Fold-regulation: Fold-change values greater than one indicate an increase in transcript abundance, relative to that of LV tissue from sham MK2^+/+^ mice, and the fold-regulation is equal to the fold-change. Where the transcript abundance is less than that of LV tissue from sham MK2^+/+^ mice, the fold-change is less than one and the fold-regulation is the negative inverse of the fold-change. *P*-values are indicated in parentheses. *N* = 3 or 4 (MK2^+/+^ sham).
