## Supplementary Table 4 for "MK2 deficiency decreases mortality during the inflammatory phase after myocardial infarction in mice"

**Supplementary Table 4. RT2 profiler PCR array analysis of chemokine mRNA in mouse left ventricular tissue 5 days post-MI.**

| Chemokine | | MK2^+/+^ | | MK2^-/-^ | | |
| --- | --- | --- | --- | --- | --- | --- |
| Symbol | Official full name | Infarct tissues | Healthy tissues | Sham | Infarct tissues | Healthy tissues |
| *Ccl1* | chemokine (C-C motif) ligand 1 | -1.64 (0.261) | -1.45 (0.560) | -2.62 (0.173) | -2.83 (0.188) | -1.82 (0.396) |
| ***Ccl2*** | **chemokine (C-C motif) ligand 2** | 1.87 (0.220) | 1.98 (0.164) | -1.07 (0.846) | **2.39 (0.047)** | 3.96 (0.108) |
| ***Ccl3*** | **chemokine (C-C motif) ligand 3** | 2.97 (0.096) | 2.40 (0.079) | -1.08 (0.761) | **4.21 (0.010)** | 3.09 (0.200) |
| ***Ccl4*** | **chemokine (C-C motif) ligand 4** | 2.75 (0.138) | 2.21 (0.144) | -1.23 (0.371) | **5.91 (0.007)** | 4.13 (0.065) |
| ***Ccl5*** | **chemokine (C-C motif) ligand 5** | 2.32 (0.110) | 1.72 (0.085) | -1.59 (0.489) | **2.54 (0.023)** | 1.41 (0.226) |
| *Ccl7* | chemokine (C-C motif) ligand 7 | 3.10 (0.098) | 1.67 (0.341) | 1.01 (0.845) | 2.41 (0.127) | 4.25 (0.204) |
| ***Ccl11*** | **chemokine (C-C motif) ligand 11** | **-5.26 (0.002)** | **-2.31 (0.022)** | 1.29 (0.392) | **-27.8 (0.001)** | -3.37 (0.098) |
| *Ccl12* | chemokine (C-C motif) ligand 12 | 1.89 (0.116) | 1.54 (0.310) | -1.63 (0.866) | 1.01 (0.810) | 1.16 (0.477) |
| *Ccl17* | chemokine (C-C motif) ligand 17 | 3.50 (0.202) | -1.11 (0.500) | 1.37 (0.786) | 3.82 (0.184) | 2.40 (0.453) |
| *Ccl19* | chemokine (C-C motif) ligand 19 | -1.38 (0.723) | -1.22 (0.468) | 1.60 (0.198) | -1.16 (0.625) | 1.91 (0.396) |
| *Ccl20* | chemokine (C-C motif) ligand 20 | 1.38 (0.818) | 1.21 (0.906) | 1.06 (0.991) | -1.35 (0.465) | 1.62 (0.616) |
| *Ccl22* | chemokine (C-C motif) ligand 22 | -2.68 (0.158) | -4.62 (0.152) | -1.84 (0.445) | -7.36 (0.133) | -6.02 (0.133) |
| *Ccl24* | chemokine (C-C motif) ligand 22 | -1.25 (0.740) | -2.74 (0.179) | 2.41 (0.263) | -2.21 (0.295) | -1.77 (0.588) |
| *Cx3cl1* | chemokine (C-X3-C motif) ligand 1 | -1.34 (0.737) | -1.14 (0.593) | -1.06 (0.939) | 1.52 (0.265) | 1.02 (0.956) |
| *Cxcl1* | chemokine (C-X-C motif) ligand 1 | -2.65 (0.184) | -2.16 (0.716) | -1.01 (0.929) | -1.47 (0.999) | -2.76 (0.604) |
| *Cxcl3* | chemokine (C-X-C motif) ligand 3 | 2.39 (0.269) | 1.44 (0.878) | -1.96 (0.281) | 2.04 (0.434) | 3.78 (0.373) |
| *Cxcl5* | chemokine (C-X-C motif) ligand 5 | 1.71 (0.691) | -3.96 (0.273) | -1.32 (0.708) | -1.09 (0.960) | -1.68 (0.654) |
| *Cxcl9* | chemokine (C-X-C motif) ligand 9 | -2.93 (0.130) | -4.17 (0.158) | -1.52 (0.677) | -3.65 (0.266) | -1.03 (0.707) |
| *Cxcl10* | chemokine (C-X-C motif) ligand 10 | 2.85 (0.261) | 2.95 (0.174) | 1.09 (0.945) | 3.81 (0.074) | 2.97 (0.080) |
| *Cxcl11* | chemokine (C-X-C motif) ligand 11 | 1.35 (0.813) | 1.42 (0.868) | -1.20 (0.579) | 1.11 (0.821) | 1.36 (0.949) |
| ***Cxcl12*** | **chemokine (C-X-C motif) ligand 12** | -1.09 (0.697) | -1.16 (0.528) | -1.02 (0.904) | **-1.49 (0.025)** | -1.27 (0.070) |
| *Cxcl13* | chemokine (C-X-C motif) ligand 13 | 3.14 (0.276) | 2.37 (0.282) | -1.93 (0.936) | 1.13 (0.598) | 1.41 (0.362) |
| *Cxcl16* | chemokine (C-X-C motif) ligand 16 | 2.99 (0.071) | 1.79 (0.133) | 1.21 (0.698) | 2.37 (0.133) | 1.87 (0.118) |
| ***Pf4*** | **platelet factor 4** | 3.16 (0.113) | 2.01 (0.180) | 1.02 (0.784) | **2.58 (0.035)** | 1.77 (0.100) |
| *Ppbp* | pro-platelet basic protein | 1.66 (0.447) | 1.04 (0.670) | 1.03 (0.643) | 1.02 (0.893) | 1.46 (0.181) |
| *Xcl1* | chemokine (C motif) ligand 1 | -1.07 (0.643) | -4.39 (0.178) | -1.20 (0.849) | -2.15 (0.389) | 1.09 (0.868) |

Data shown are expressed as the fold-regulation in transcript abundance relative to LV tissue from sham MK2^+/+^ mice. Fold-regulation: Fold-change values greater than one indicate an increase in transcript abundance, relative to that of LV tissue from sham MK2^+/+^ mice, and the fold-regulation is equal to the fold-change. Where the transcript abundance is less than that of LV tissue from sham MK2^+/+^ mice, the fold-change is less than one and the fold-regulation is the negative inverse of the fold-change. *P*-values are indicated in parentheses. *N* = 3 or 4 (MK2^+/+^ infarct tissue).
