## Supplementary Table 5 for "MK2 deficiency decreases mortality during the inflammatory phase after myocardial infarction in mice"

**Supplementary Table 5. RT2 profiler PCR array analysis of growth factor mRNA in mouse left ventricular tissue 3 days post-MI.**

| Growth factor | | MK2^+/+^ | | MK2^-/-^ | | |
| --- | --- | --- | --- | --- | --- | --- |
| Symbol | Official full name | Infarct tissues | Healthy tissues | Sham | Infarct tissues | Healthy tissues |
| ***Bmp2*** | **bone morphogenetic protein 2** | **9.56 (0.001)** | 2.74 (0.149) | -1.19 (0.444) | 5.35 (0.060) | 1.44 (0.963) |
| *Bmp4* | bone morphogenetic protein 4 | 1.09 (0.973) | 1.29 (0.660) | -1.05 (0.898) | 1.27 (0.661) | 1.45 (0.444) |
| ***Bmp6*** | **bone morphogenetic protein 6** | -2.17 (0.051) | **-2.57 (0.033)** | -1.75 (0.103) | **-2.64 (0.026)** | **-2.56 (0.037)** |
| ***Bmp7*** | **bone morphogenetic protein 7** | **-6.10 (0.011)** | **-3.31 (0.020)** | -1.31 (0.448) | **-10.8 (0.005)** | -3.10 (0.319) |
| *Cntf* | ciliary neurotrophic factor | 1.02 (0.858) | -1.48 (0.333) | -1.71 (0.150) | -1.37 (0.279) | 1.15 (0.451) |
| ***Csf1*** | **colony stimulating factor 1 (macrophage)** | -1.22 (0.254) | -1.61 (0.115) | **-2.14 (0.038)** | **-1.92 (0.026)** | **-2.22 (0.015)** |
| ***Csf2*** | **colony stimulating factor 2 (granulocyte-macrophage)** | 3.85 (0.215) | **4.26 (0.037)** | 2.13 (0.210) | **12.7 (0.004)** | **4.10 (0.024)** |
| *Csf3* | colony stimulating factor 3 (granulocyte) | 4.21 (0.100) | 2.12 (0.192) | 1.87 (0.178) | 2.40 (0.199) | 3.16 (0.114) |
| ***Gpi1*** | **glucose phosphate isomerase 1** | **-1.99 (0.004)** | **-1.75 (0.007)** | -1.26 (0.255) | **-2.28 (0.003)** | **-1.61 (0.011)** |
| *Lif* | leukemia inhibitory factor | 4.18 (0.058) | 1.71 (0.264) | 1.04 (0.489) | 2.53 (0.170) | -1.85 (0.939) |
| *Mstn* | myostatin | -1.25 (0.899) | -1.30 (0.486) | -1.24 (0.847) | -1.51 (0.437) | 1.74 (0.369) |
| *Nodal* | nodal | -1.35 (0.388) | -1.11 (0.562) | 1.28 (0.512) | 1.13 (0.778) | 2.20 (0.239) |
| ***Osm*** | **oncostatin M** | 13.6 (0.111) | 6.10 (0.081) | 1.15 (0.448) | **11.5 (0.003)** | -1.13 (0.976) |
| *Thpo* | thrombopoietin | -1.91 (0.524) | -2.27 (0.314) | -1.27 (0.737) | -2.25 (0.335) | -1.62 (0.831) |
| ***Vegfa*** | **vascular endothelial growth factor A** | **-3.35 (0.016)** | **-2.62 (0.017)** | -2.28 (0.071) | **-3.50 (0.012)** | **-2.69 (0.017)** |

Data shown are expressed as the fold-regulation in transcript abundance relative to LV tissue from sham MK2^+/+^ mice. Fold-regulation: Fold-change values greater than one indicate an increase in transcript abundance, relative to that of LV tissue from sham MK2^+/+^ mice, and the fold-regulation is equal to the fold-change. Where the transcript abundance is less than that of LV tissue from sham MK2^+/+^ mice, the fold-change is less than one and the fold-regulation is the negative inverse of the fold-change. *P*-values are indicated in parentheses.  *N* = 3 or 4 (MK2^+/+^ sham).
