## Supplementary Table 6 for "MK2 deficiency decreases mortality during the inflammatory phase after myocardial infarction in mice"

**Supplementary Table 6. RT2 profiler PCR array analysis of growth factor mRNA in mouse left ventricular tissue 5 days post-MI.**

| Growth factor | | MK2^+/+^ | | MK2^-/-^ | | |
| --- | --- | --- | --- | --- | --- | --- |
| Symbol | Official full name | Infarct tissues | Healthy tissues | Sham | Infarct tissues | Healthy tissues |
| *Bmp2* | bone morphogenetic protein 2 | 1.38 (0.764) | 1.46 (0.831) | -2.31 (0.582) | 1.73 (0.629) | -1.27 (0.495) |
| *Bmp4* | bone morphogenetic protein 4 | 1.47 (0.126) | 1.44 (0.201) | 1.51 (0.260) | -1.52 (0.961) | 1.18 (0.483) |
| ***Bmp6*** | **bone morphogenetic protein 6** | **-2.30 (0.015)** | -1.29 (0.424) | 1.55 (0.068) | **-2.10 (0.027)** | -1.63 (0.111) |
| ***Bmp7*** | **bone morphogenetic protein 7** | **-6.10 (0.0004)** | **-2.76 (0.003)** | -1.25 (0.663) | **-4.17 (0.001)** | -2.12 (0.381) |
| *Cntf* | ciliary neurotrophic factor | 1.44 (0.255) | 1.11 (0.434) | -1.05 (0.879) | -1.03 (0.911) | 1.25 (0.376) |
| ***Csf1*** | **colony stimulating factor 1 (macrophage)** | -1.67 (0.060) | -1.23 (0.073) | -1.22 (0.185) | **-1.44 (0.032)** | -1.40 (0.074) |
| *Csf2* | colony stimulating factor 2 (granulocyte-macrophage) | -3.32 (0.288) | -10.10 (0.374) | 1.89 (0.428) | -2.02 (0.380) | -4.35 (0.377) |
| *Csf3* | colony stimulating factor 3 (granulocyte) | -1.15 (0.573) | -1.31 (0.373) | -1.02 (0.994) | -1.12 (0.834) | 2.36 (0.400) |
| ***Gpi1*** | **glucose phosphate isomerase 1** | **-2.29 (0.001)** | -1.51 (0.062) | 1.08 (0.484) | **-2.43 (0.002)** | **-1.48 (0.009)** |
| *Lif* | leukemia inhibitory factor | 2.33 (0.328) | 1.22 (0.661) | 3.15 (0.051) | 4.06 (0.124) | 5.29 (0.093) |
| *Mstn* | myostatin | -1.24 (0.412) | -1.49 (0.299) | 2.89 (0.109) | -1.72 (0.335) | -1.05 (0.814) |
| *Nodal* | nodal | -3.14 (0.059) | -3.08 (0.101) | -1.23 (0.808) | -5.85 (0.057) | 1.21 (0.731) |
| ***Osm*** | **oncostatin M** | **8.53 (0.037)** | 3.58 (0.182) | 2.25 (0.206) | **6.05 (0.010)** | 1.24 (0.608) |
| *Thpo* | thrombopoietin | -1.16 (0.753) | 1.59 (0.418) | 1.04 (0.890) | 1.28 (0.709) | 1.26 (0.629) |
| ***Vegfa*** | **vascular endothelial growth factor A** | **-4.86 (0.032)** | -1.32 (0.397) | 1.08 (0.901) | -3.82 (0.072) | -1.26 (0.506) |

Data shown are expressed as the fold-regulation in transcript abundance relative to LV tissue from sham MK2^+/+^ mice. Fold-regulation: Fold-change values greater than one indicate an increase in transcript abundance, relative to that of LV tissue from sham MK2^+/+^ mice, and the fold-regulation is equal to the fold-change. Where the transcript abundance is less than that of LV tissue from sham MK2^+/+^ mice, the fold-change is less than one and the fold-regulation is the negative inverse of the fold-change. *P*-values are indicated in parentheses. *N* = 3 or 4 (MK2^+/+^ infarct tissue).
