## Supplementary Table 7 for "MK2 deficiency decreases mortality during the inflammatory phase after myocardial infarction in mice"

**Supplementary Table 7. RT2 profiler PCR array analysis of interferon mRNA in mouse left ventricular tissue 3 days post-MI.**

| Interferon | | MK2^+/+^ | | MK2^-/-^ | | |
| --- | --- | --- | --- | --- | --- | --- |
| Symbol | Official full name | Infarct tissues | Healthy tissues | Sham | Infarct tissues | Healthy tissues |
| ***Ifna2*** | **interferon alpha 2** | 1.45 (0.238) | 4.44 (0.109) | 3.27 (0.175) | **14.5 (0.005)** | 7.01 (0.194) |
| *Ifng* | interferon gamma | -1.35 (0.388) | -1.08 (0.606) | -1.34 (0.344) | 1.13 (0.778) | 1.90 (0.125) |

Data shown are expressed as the fold-regulation in transcript abundance relative to LV tissue from sham MK2^+/+^ mice. Fold-regulation: Fold-change values greater than one indicate an increase in transcript abundance, relative to that of LV tissue from sham MK2^+/+^ mice, and the fold-regulation is equal to the fold-change. Where the transcript abundance is less than that of LV tissue from sham MK2^+/+^ mice, the fold-change is less than one and the fold-regulation is the negative inverse of the fold-change. *P*-values are indicated in parentheses.  *N* = 3 or 4 (MK2^+/+^ sham).
