## Supplementary Table 8 for "MK2 deficiency decreases mortality during the inflammatory phase after myocardial infarction in mice"

**Supplementary Table 8. RT2 profiler PCR array analysis of interferon mRNA in mouse left ventricular tissue 5 days post-MI.**

| Interferon | | MK2^+/+^ | | MK2^-/-^ | | |
| --- | --- | --- | --- | --- | --- | --- |
| Symbol | Official full name | Infarct tissues | Healthy tissues | Sham | Infarct tissues | Healthy tissues |
| *Ifna2* | interferon alpha 2 | 1.02 (0.452) | -1.22 (0.299) | 1.48 (0.511) | -1.54 (0.311) | -1.21 (0.715) |
| *Ifng* | interferon gamma | 2.27 (0.340) | 1.21 (0.906) | 1.06 (0.991) | 1.52 (0.562) | 1.62 (0.616) |

Data shown are expressed as the fold-regulation in transcript abundance relative to LV tissue from sham MK2^+/+^ mice. Fold-regulation: Fold-change values greater than one indicate an increase in transcript abundance, relative to that of LV tissue from sham MK2^+/+^ mice, and the fold-regulation is equal to the fold-change. Where the transcript abundance is less than that of LV tissue from sham MK2^+/+^ mice, the fold-change is less than one and the fold-regulation is the negative inverse of the fold-change. *P*-values are indicated in parentheses. *N* = 3 or 4 (MK2^+/+^ infarct tissue).
