## Supplementary Table 9 for "MK2 deficiency decreases mortality during the inflammatory phase after myocardial infarction in mice"

**Supplementary Table 9. RT2 profiler PCR array analysis of interleukin mRNA in mouse left ventricular tissue 3 days post-MI.**

| Interleukin | | MK2^+/+^ | | MK2^-/-^ | | |
| --- | --- | --- | --- | --- | --- | --- |
| Symbol | Official full name | Infarct tissues | Healthy tissues | Sham | Infarct tissues | Healthy tissues |
| *Il1a* | interleukin 1 alpha | 1.18 (0.620) | -2.10 (0.292) | -1.96 (0.427) | -1.06 (0.709) | -2.85 (0.231) |
| ***Il1b*** | **interleukin 1 beta** | 28.2 (0.144) | **12.5 (0.040)** | 6.02 (0.187) | 16.1 (0.061) | **9.32 (0.018)** |
| ***Il1rn*** | **interleukin 1 receptor antagonist** | **5.95 (0.033)** | **4.46 (0.0005)** | 1.13 (0.559) | **4.52 (0.002)** | **1.82 (0.015)** |
| *Il2* | interleukin 2 | -1.35 (0.388) | 1.30 (0.576) | 1.08 (0.793) | 1.18 (0.724) | 1.38 (0.350) |
| *Il3* | interleukin 3 | -1.35 (0.388) | 1.14 (0.862) | -1.21 (0.469) | 1.13 (0.778) | 1.38 (0.350) |
| *Il4* | interleukin 4 | -2.27 (0.215) | -1.53 (0.356) | -1.65 (0.339) | 1.37 (0.573) | -1.22 (0.831) |
| *Il5* | interleukin 5 | -2.32 (0.393) | 1.24 (0.960) | 1.25 (0.709) | 1.29 (0.867) | 2.77 (0.457) |
| ***Il6*** | **interleukin 6** | 23.7 (0.110) | 17.4 (0.055) | 1.19 (0.964) | **29.3 (0.0002)** | **9.16 (0.033)** |
| *Il7* | interleukin 7 | -2.2 (0.118) | -1.54 (0.749) | 1.26 (0.519) | -1.43 (0.395) | -1.76 (0.150) |
| *Il9* | interleukin 9 | -1.35 (0.388) | -1.11 (0.562) | -1.30 (0.360) | 1.13 (0.778) | 1.38 (0.350) |
| ***Il10*** | **interleukin 10** | 4.69 (0.149) | 5.71 (0.058) | **4.16 (0.011)** | 7.49 (0.123) | 8.88 (0.211) |
| ***Il11*** | **interleukin 11** | 5.64 (0.145) | 1.96 (0.277) | -1.37 (0.421) | **3.03 (0.035)** | 1.36 (0.383) |
| *Il12a* | interleukin 12a | -5.02 (0.122) | -7.15 (0.090) | -8.29 (0.094) | -1.47 (0.599) | -2.05 (0.317) |
| *Il12b* | interleukin 12b | -1.35 (0.388) | 1.85 (0.329) | 2.67 (0.225) | 4.22 (0.092) | 1.95 (0.103) |
| *Il13* | interleukin 13 | -1.23 (0.843) | -1.61 (0.413) | -1.21 (0.679) | -1.08 (0.640) | 1.53 (0.946) |
| ***Il15*** | **interleukin 15** | **-4.13 (0.006)** | **-2.47 (0.038)** | **-9.03 (0.003)** | **-7.57 (0.004)** | **-5.62 (0.012)** |
| ***Il16*** | **interleukin 16** | 1.33 (0.278) | -1.63 (0.164) | -1.44 (0.171) | -1.86 (0.091) | **-3.69 (0.029)** |
| *Il17a* | interleukin 17A | -1.35 (0.388) | 1.45 (0.426) | -1.04 (0.882) | 2.09 (0.266) | 1.38 (0.350) |
| *Il17f* | interleukin 17F | -3.36 (0.220) | -2.23 (0.298) | -1.45 (0.528) | -1.24 (0.654) | -1.04 0.923 |
| ***Il18*** | **interleukin 18** | 1.76 (0.446) | **5.93 (0.008)** | 1.35 (0.688) | 2.00 (0.357) | **3.82 (0.040)** |
| *Il21* | interleukin 21 | -1.35 (0.388) | -1.11 (0.562) | -1.39 (0.295) | 1.60 (0.220) | 1.38 (0.350) |
| *Il22* | interleukin 22 | -1.35 (0.388) | 1.57 (0.383) | -1.39 (0.295) | 1.13 (0.778) | 1.38 (0.350) |
| *Il23a* | interleukin 23, alpha subunit p19 | -2.13 (0.202) | 1.25 0.598 | -1.43 (0.535) | -1.01 (0.985) | -1.17 (0.549) |
| *Il24* | interleukin 24 | 1.28 (0.447) | -1.07 (0.616) | -1.39 (0.295) | 1.33 (0.481) | 1.39 (0.348) |
| ***Il27*** | **interleukin 27** | 5.75 (0.145) | 1.16 (0.806) | 1.30 (0.455) | **15.5 (0.0002)** | 5.18 (0.074) |

Data shown are expressed as the fold-regulation in transcript abundance relative to LV tissue from sham MK2^+/+^ mice. Fold-regulation: Fold-change values greater than one indicate an increase in transcript abundance, relative to that of LV tissue from sham MK2^+/+^ mice, and the fold-regulation is equal to the fold-change. Where the transcript abundance is less than that of LV tissue from sham MK2^+/+^ mice, the fold-change is less than one and the fold-regulation is the negative inverse of the fold-change. *P*-values are indicated in parentheses.  *N* = 3 or 4 (MK2^+/+^ sham).
