## Supplementary Table 10 for "MK2 deficiency decreases mortality during the inflammatory phase after myocardial infarction in mice"

**Supplementary Table 10. RT2 profiler PCR array analysis of interleukin mRNA in mouse left ventricular tissue 5 days post-MI.**

| Interleukin | | MK2^+/+^ | | MK2^-/-^ | | |
| --- | --- | --- | --- | --- | --- | --- |
| Symbol | Official full name | Infarct tissues | Healthy tissues | Sham | Infarct tissues | Healthy tissues |
| *Il1a* | interleukin 1 alpha | 2.45 (0.429) | 4.10 (0.126) | 4.27 (0.185) | 1.58 (0.569) | 21.9 (0.251) |
| *Il1b* | interleukin 1 beta | 3.93 (0.204) | 1.20 (0.720) | -1.62 (0.935) | 3.99 (0.246) | 8.11 (0.262) |
| ***Il1rn*** | **interleukin 1 receptor antagonist** | **5.08 (0.034)** | **2.88 (0.034)** | -1.15 (0.834) | **8.64 (0.002)** | 3.23 (0.142) |
| *Il2* | interleukin 2 | 1.38 (0.818) | 1.21 (0.906) | 1.07 (0.980) | -1.35 (0.465) | 1.62 (0.616) |
| *Il3* | interleukin 3 | 1.38 (0.818) | 1.21 (0.906) | 1.06 (0.991) | -1.35 (0.465) | 1.62 (0.616) |
| *Il4* | interleukin 4 | 1.56 (0.605) | 1.21 (0.906) | 1.70 (0.478) | 2.30 (0.288) | 1.62 (0.616) |
| ***Il5*** | **interleukin 5** | -1.85 (0.491) | **-3.75 (0.031)** | -1.56 (0.921) | **-3.94 (0.035)** | -1.32 (0.884) |
| *Il6* | interleukin 6 | 5.64 (0.113) | 4.00 (0.173) | 1.35 (0.513) | 15.9 (0.156) | 13.2 (0.328) |
| *Il7* | interleukin 7 | -1.63 (0.372) | -1.55 (0.598) | 2.15 (0.223) | -1.54 (0.343) | 1.66 (0.416) |
| *Il9* | interleukin 9 | 1.16 (0.956) | 1.02 (0.754) | -1.12 (0.891) | -1.61 (0.340) | 1.36 (0.687) |
| ***Il10*** | **interleukin 10** | 1.78 (0.342) | 2.26 (0.225) | 3.27 (0.068) | **8.38 (0.026)** | 6.07 (0.127) |
| *Il11* | interleukin 11 | 5.38 (0.203) | 3.59 (0.279) | 1.32 (0.628) | 1.18 (0.746) | 2.24 (0.243) |
| *Il12a* | interleukin 12a | 1.89 (0.302) | 1.48 (0.530) | 1.76 (0.318) | -2.12 (0.449) | 1.71 (0.377) |
| *Il12b* | interleukin 12b | -1.36 (0.373) | -1.41 (0.360) | 1.31 (0.540) | -1.98 (0.271) | 1.56 (0.498) |
| *Il13* | interleukin 13 | 1.16 (0.747) | 1.48 (0.730) | 1.64 (0.480) | -1.53 (0.387) | 2.41 (0.405) |
| ***Il15*** | **interleukin 15** | **-7.52 (0.001)** | **-3.59 (0.012)** | **-3.14 (0.039)** | **-6.23 (0.006)** | **-8.20 (0.006)** |
| ***Il16*** | **interleukin 16** | -2.23 (0.167) | -1.46 (0.244) | **1.96 (0.009)** | -1.19 (0.536) | -4.63 (0.194) |
| *Il17a* | interleukin 17A | -1.21 (0.495) | -1.38 (0.335) | -1.10 (0.923) | -2.25 (0.146) | -1.03 (0.924) |
| *Il17f* | interleukin 17F | -1.94 (0.296) | -2.61 (0.233) | 1.66 (0.524) | -4.26 (0.174) | -1.77 (0.507) |
| *Il18* | interleukin 18 | 1.95 (0.224) | 1.14 (0.990) | -1.17 (0.952) | 1.16 (0.862) | -1.96 (0.828) |
| *Il21* | interleukin 21 | 1.38 (0.818) | 1.21 (0.906) | 1.06 (0.991) | -1.35 (0.465) | 1.62 (0.616) |
| *Il22* | interleukin 22 | 1.38 (0.818) | 1.21 (0.906) | 1.43 (0.690) | -1.35 (0.465) | 1.62 (0.616) |
| ***Il23a*** | **interleukin 23, alpha subunit p19** | **-2.61 (0.025)** | -1.45 (0.428) | -1.01 (0.801) | -1.89 (0.353) | -1.77 (0.750) |
| *Il24* | interleukin 24 | 1.38 (0.818) | 1.21 (0.906) | 1.06 (0.991) | -1.17 (0.580) | 1.62 (0.616) |
| *Il27* | interleukin 27 | 5.96 (0.228) | 2.32 (0.268) | -1.16 (0.744) | 4.60 (0.064) | -1.17 (0.700) |

Data shown are expressed as the fold-regulation in transcript abundance relative to LV tissue from sham MK2^+/+^ mice. Fold-regulation: Fold-change values greater than one indicate an increase in transcript abundance, relative to that of LV tissue from sham MK2^+/+^ mice, and the fold-regulation is equal to the fold-change. Where the transcript abundance is less than that of LV tissue from sham MK2^+/+^ mice, the fold-change is less than one and the fold-regulation is the negative inverse of the fold-change. *P*-values are indicated in parentheses. *N* = 3 or 4 (MK2^+/+^ infarct tissue).
