## Supplementary Table 11 for "MK2 deficiency decreases mortality during the inflammatory phase after myocardial infarction in mice"

**Supplementary Table 11. RT2 profiler PCR array analysis of other cytokine mRNA in mouse left ventricular tissue 3 days post-MI.**

| Other cytokine | | MK2^+/+^ | | MK2^-/-^ | | |
| --- | --- | --- | --- | --- | --- | --- |
| Symbol | Official full name | Infarct tissues | Healthy tissues | Sham | Infarct tissues | Healthy tissues |
| *Adipoq* | adiponectin, C1Q and collagen domain containing | -4.23 (0.299) | 1.63 (0.466) | -1.35 (0.660) | -2.78 (0.350) | -1.52 (0.462) |
| ***Ctf1*** | **cardiotrophin 1** | **-4.93 (0.008)** | **-3.31 (0.016)** | -1.51 (0.254) | **-9.31 (0.005)** | -3.56 (0.053) |
| *Hc* | hemolytic complement | 2.13 (0.434) | -1.29 (0.434) | -1.50 (0.467) | 1.40 (0.961) | 1.13 (0.652) |
| *Mif* | macrophage migration inhibitory factor (glycosylation-inhibiting factor) | 1.00 (0.970) | 1.17 (0.440) | 1.14 (0.507) | 1.22 (0.294) | 1.60 (0.169) |
| ***Spp1*** | **secreted phosphoprotein 1** | **32.6 (0.004)** | **16.5 (0.00001)** | 3.90 (0.161) | **25.2 (0.0001)** | **18.6 (0.008)** |
| ***Tgfb2*** | **transforming growth factor, beta 2** | **2.04 (0.023)** | **1.91 (0.002)** | -1.43 (0.071) | **3.01 (0.040)** | 1.01 (0.827) |

Data shown are expressed as the fold-regulation in transcript abundance relative to LV tissue from sham MK2^+/+^ mice. Fold-regulation: Fold-change values greater than one indicate an increase in transcript abundance, relative to that of LV tissue from sham MK2^+/+^ mice, and the fold-regulation is equal to the fold-change. Where the transcript abundance is less than that of LV tissue from sham MK2^+/+^ mice, the fold-change is less than one and the fold-regulation is the negative inverse of the fold-change. *P*-values are indicated in parentheses.  *N* = 3 or 4 (MK2^+/+^ sham).
