## Supplementary Table 12 for "MK2 deficiency decreases mortality during the inflammatory phase after myocardial infarction in mice"

**Supplementary Table 12. RT2 profiler PCR array analysis of other cytokine mRNA in mouse left ventricular tissue 5 days post-MI.**

| Other cytokines | | MK2^+/+^ | | MK2^-/-^ | | |
| --- | --- | --- | --- | --- | --- | --- |
| Symbol | Official full name | Infarct tissues | Healthy tissues | Sham | Infarct tissues | Healthy tissues |
| *Adipoq* | adiponectin, C1Q and collagen domain containing | 1.62 (0.434) | 1.07 (0.606) | 1.29 (0.551) | -4.49 (0.060) | -1.61 (0.382) |
| ***Ctf1*** | **cardiotrophin 1** | **-11.7 (0.007)** | **-5.61 (0.024)** | -1.09 (0.776) | **-4.31 (0.029)** | -4.30 (0.054) |
| *Hc* | hemolytic complement | -1.03 (0.652) | 2.91 (0.267) | 1.04 (0.637) | 2.51 (0.263) | 1.72 (0.557) |
| *Mif* | macrophage migration inhibitory factor (glycosylation-inhibiting factor) | 1.12 (0.906) | 1.03 (0.941) | 1.27 (0.625) | -1.43 (0.343) | 1.19 (0.829) |
| ***Spp1*** | **secreted phosphoprotein 1** | **38.2 (0.009)** | **7.52 (0.006)** | 1.15 (0.847) | **30.5 (0.036)** | 8.88 (0.126) |
| ***Tgfb2*** | **transforming growth factor, beta 2** | **2.32 (0.049)** | 1.56 (0.059) | **1.11 (0.031)** | **1.82 (0.031)** | -1.20 (0.659) |

Data shown are expressed as the fold-regulation in transcript abundance relative to LV tissue from sham MK2^+/+^ mice. Fold-regulation: Fold-change values greater than one indicate an increase in transcript abundance, relative to that of LV tissue from sham MK2^+/+^ mice, and the fold-regulation is equal to the fold-change. Where the transcript abundance is less than that of LV tissue from sham MK2^+/+^ mice, the fold-change is less than one and the fold-regulation is the negative inverse of the fold-change. *P*-values are indicated in parentheses. *N* = 3 or 4 (MK2^+/+^ infarct tissue).
