## Supplementary Table 13 for "MK2 deficiency decreases mortality during the inflammatory phase after myocardial infarction in mice"

**Supplementary Table 13. RT2 profiler PCR array analysis of TNF receptor superfamily member mRNA in mouse left ventricular tissue 3 days post-MI.**

| TNF receptor superfamily member | | MK2^+/+^ | | MK2^-/-^ | | |
| --- | --- | --- | --- | --- | --- | --- |
| Symbol | Official full name | Infarct tissues | Healthy tissues | Sham | Infarct tissues | Healthy tissues |
| *Cd40lg* | CD40 ligand | -1.74 (0.390) | -1.43 (0.443) | 1.03 (0.957) | 1.26 (0.875) | 1.69 (0.775) |
| *Cd70* | CD70 antigen | 1.26 (0.555) | -1.11 (0.562) | -1.39 (0.295) | 1.13 (0.778) | 1.38 (0.350) |
| *Fasl* | Fas ligand (TNF superfamily, member 6) | -1.69 (0.182) | -1.39 (0.250) | -1.75 (0.138) | -1.11 (0.747) | 1.87 (0.298) |
| *Lta* | lymphotoxin A | -1.35 (0.388) | 1.52 (0.393) | 2.66 (0.105) | 1.47 (0.401) | 1.47 (0.317) |
| ***Ltb*** | **lymphotoxin B** | 2.08 (0.169) | -1.72 (0.806) | **2.32 (0.009)** | -1.00 (0.675) | -2.06 (0.101) |
| *Tnf* | tumor necrosis factor | 1.33 (0.965) | 1.56 (0.901) | -1.05 (0.619) | 1.84 (0.895) | -1.38 (0.488) |
| ***Tnfrsf11b*** | **tumor necrosis factor receptor superfamily, member 11b (osteoprotegerin)** | **8.64 (0.036)** | **11.9 (0.004)** | 1.83 (0.790) | **10.3 (0.002)** | 7.03 (0.168) |
| *Tnfsf10* | tumor necrosis factor (ligand) superfamily, member 10 | -1.08 (0.688) | -1.04 (0.812) | 1.18 (0.601) | 1.13 (0.720) | -1.96 (0.678) |
| ***Tnfsf11*** | **tumor necrosis factor (ligand) superfamily, member 11** | 30.8 (0.070) | **14.9 (0.001)** | 4.00 (0.080) | **21.1 (0.021)** | 4.87 (0.052) |
| ***Tnfsf13b*** | **tumor necrosis factor (ligand) superfamily, member 13b** | -2.18 (0.080) | -2.00 (0.124) | -2.12 (0.160) | **-8.25 (0.028)** | -2.40 (0.071) |

Data shown are expressed as the fold-regulation in transcript abundance relative to LV tissue from sham MK2^+/+^ mice. Fold-regulation: Fold-change values greater than one indicate an increase in transcript abundance, relative to that of LV tissue from sham MK2^+/+^ mice, and the fold-regulation is equal to the fold-change. Where the transcript abundance is less than that of LV tissue from sham MK2^+/+^ mice, the fold-change is less than one and the fold-regulation is the negative inverse of the fold-change. *P*-values are indicated in parentheses. *N* = 3 or 4 (MK2^+/+^ sham).
