## Supplementary Table 14 for "MK2 deficiency decreases mortality during the inflammatory phase after myocardial infarction in mice"

**Supplementary Table 14. RT2 profiler PCR array analysis of TNF receptor superfamily member mRNA in mouse left ventricular tissue 5 days post-MI.**

| TNF receptors superfamily member | | MK2^+/+^ | | MK2^-/-^ | | |
| --- | --- | --- | --- | --- | --- | --- |
| Symbol | Official full name | Infarct tissues | Healthy tissues | Sham | Infarct tissues | Healthy tissues |
| ***Cd40lg*** | **CD40 ligand** | 1.90 (0.288) | **-1.79 (0.001)** | -2.42 (0.133) | -1.10 (0.686) | -1.06 (0.585) |
| *Cd70* | CD70 antigen | 1.72 (0.510) | 1.21 (0.906) | 1.06 (0.991) | -1.35 (0.465) | 1.62 (0.616) |
| *Fasl* | Fas ligand (TNF superfamily, member 6) | 2.03 (0.365) | 1.21 (0.906) | 1.06 (0.991) | -1.35 (0.465) | 1.62 (0.616) |
| *Lta* | lymphotoxin A | 2.49 (0.443) | 1.16 (0.875) | 1.01 (0.985) | 1.46 (0.565) | 2.11 (0.344) |
| *Ltb* | lymphotoxin B | 1.78 (0.513) | 1.63 (0.721) | 1.31 (0.779) | 2.02 (0.392) | 1.07 (0.979) |
| *Tnf* | tumor necrosis factor | -1.07 (0.823) | -1.05 (0.658) | -1.26 (0.887) | -1.41 (0.315) | -2.69 (0.520) |
| ***Tnfrsf11b*** | **tumor necrosis factor receptor superfamily, member 11b (osteoprotegerin)** | **2.82 (0.011)** | 1.43 (0.390) | -1.41 (0.294) | **2.83 (0.009)** | **2.65 (0.029)** |
| *Tnfsf10* | tumor necrosis factor (ligand) superfamily, member 10 | -2.08 (0.082) | -1.28 (0.438) | 2.07 (0.075) | -1.80 (0.174) | 1.17 (0.882) |
| ***Tnfsf11*** | **tumor necrosis factor (ligand) superfamily, member 11** | **4.35 (0.046)** | 1.65 (0.452) | -1.03 (0.887) | 3.07 (0.234) | 2.89 (0.356) |
| ***Tnfsf13b*** | **tumor necrosis factor (ligand) superfamily, member 13b** | **-1.55 (0.020)** | 1.33 (0.350) | 1.50 (0.128) | **-1.80 (0.017)** | -1.58 (0.647) |

Data shown are expressed as the fold-regulation in transcript abundance relative to LV tissue from sham MK2^+/+^ mice. Fold-regulation: Fold-change values greater than one indicate an increase in transcript abundance, relative to that of LV tissue from sham MK2^+/+^ mice, and the fold-regulation is equal to the fold-change. Where the transcript abundance is less than that of LV tissue from sham MK2^+/+^ mice, the fold-change is less than one and the fold-regulation is the negative inverse of the fold-change. *P*-values are indicated in parentheses. *N* = 3 or 4 (MK2^+/+^ infarct tissue).
