## Supplementary Table 15 for "MK2 deficiency decreases mortality during the inflammatory phase after myocardial infarction in mice"

**Supplementary Table 15. MK2 dependence of cytokine transcripts induced by lipopolysaccharide differs 3 days post-MI.**

| 3 days | | | | | | |
| --- | --- | --- | --- | --- | --- | --- |
| Cytokine | | MK2^+/+^ | | MK2^-/-^ | | |
| Symbol | Official full name | Healthy tissue (n = 3) | Infarct tissue (n = 3) | Sham  (n = 3) | Healthy tissue (n = 3) | Infarct tissue (n = 3) |
| *Ccl3* | chemokine (C-C motif) ligand 3 | **3.67 (0.002)** | 5.12 (0.079) | 1.07 (0.927) | 1.50 (0.212) | 3.83 (0.053) |
| *Ccl4* | chemokine (C-C motif) ligand 4 | **2.81 (0.001)** | **4.61 (0.011)** | -1.44 (0.524) | 1.73 (0.175) | 2.84 (0.067) |
| *Ccl19* | chemokine (C-C motif) ligand 19 | -1.49 (0.318) | **-2.34 (0.032)** | -1.59 (0.129) | 1.34 (0.353) | **-2.03 (0.047)** |
| *Ccl20* | chemokine (C-C motif) ligand 20 | -1.11 (0.562) | -1.35 (0.388) | -1.39 (0.295) | 1.38 (0.350) | 1.85 (0.151) |
| *Cxcl3* | chemokine (C-X-C motif) ligand 3 | **27.3 (0.021)** | 74.3 (0.163) | 2.55 (0.194) | **5.57 (0.044)** | **20.2 (0.023)** |
| *Il1b* | interleukin 1 beta | **12.5 (0.040)** | 28.2 (0.144) | 6.02 (0.187) | **9.32 (0.018)** | 16.1 (0.061) |
| *Il6* | interleukin 6 | 17.4 (0.055) | 23.7 (0.110) | 1.19 (0.964) | **9.16 (0.033)** | **29.3 (0.0002)** |
| *Il10* | interleukin 10 | 5.71 (0.058) | 4.69 (0.149) | **4.16 (0.011)** | 8.88 (0.211) | 7.49 (0.123) |
| *Il12a* | interleukin 12a | -7.15 (0.090) | -5.02 (0.122) | -8.29 (0.094) | -2.05 (0.317) | -1.47 (0.599) |
| *Il12b* | interleukin 12b | 1.85 (0.329) | -1.35 (0.388) | 2.67 (0.225) | 1.95 (0.103) | 4.22 (0.092) |
| *Il15* | interleukin 15 | **-2.47 (0.038)** | **-4.13 (0.006)** | **-9.03 (0.003)** | **-5.62 (0.012)** | **-7.57 (0.004)** |
| *Ifng* | interferon gamma | -1.08 (0.606) | -1.35 (0.388) | -1.34 (0.344) | 1.90 (0.125) | 1.13 (0.778) |
| *Csf2* | colony stimulating factor 2 (granulocyte-macrophage) | **4.26 (0.037)** | 3.85 (0.215) | 2.13 (0.210) | **4.10 (0.024)** | **12.7 (0.004)** |
| *Vegfa* | vascular endothelial growth factor A | **-2.62 (0.017)** | **-3.35 (0.016)** | -2.28 (0.071) | **-2.69 (0.017)** | **-3.50 (0.012)** |
| *Tnf* | tumor necrosis factor | 1.56 (0.901) | 1.33 (0.965) | -1.05 (0.619) | -1.38 (0.488) | 1.84 (0.895) |
| *Tgfb2* | transforming growth factor, beta 2 | **1.91 (0.002)** | **2.04 (0.023)** | -1.43 (0.071) | 1.01 (0.827) | **3.01 (0.040)** |

Data shown are expressed as the fold-regulation in transcript abundance relative to 3-day sham MK2^+/+^. Fold-regulation: fold-change values greater than one indicate an increase in transcript abundance, relative to that of sham MK2^+/+^ hearts, and the fold-regulation is equal to the fold-change. Where the transcript abundance is less than that of sham MK2^+/+^ hearts, the fold-change is less than one and the fold-regulation is the negative inverse of the fold-change. *P*-values are indicated in parentheses.
