## Supplementary Table 16 for "MK2 deficiency decreases mortality during the inflammatory phase after myocardial infarction in mice"

**Supplementary Table 16. MK2 dependence of cytokine transcripts induced by lipopolysaccharide differs 5 days post-MI.**

| 5 Days | | | | | | |
| --- | --- | --- | --- | --- | --- | --- |
| Cytokine | | MK2^+/+^ | | MK2^-/-^ | | |
| Symbol | Official full name | Healthy tissues  (n = 3) | Infarct tissues (n = 4) | Sham (n = 3) | Healthy tissues (n = 3) | Infarct tissues (n = 3) |
| *Ccl3* | chemokine (C-C motif) ligand 3 | 2.40 (0.079) | 2.97 (0.096) | -1.08 (0.761) | 3.09 (0.200) | **4.21 (0.010)** |
| *Ccl4* | chemokine (C-C motif) ligand 4 | 2.21 (0.144) | 2.75 (0.138) | -1.23 (0.371) | 4.13 (0.065) | **5.91 (0.007)** |
| *Ccl19* | chemokine (C-C motif) ligand 19 | -1.22 (0.468) | -1.38 (0.723) | 1.60 (0.198) | 1.91 (0.396) | -1.16 (0.625) |
| *Ccl20* | chemokine (C-C motif) ligand 20 | 1.21 (0.906) | 1.38 (0.818) | 1.06 (0.991) | 1.62 (0.616) | -1.35 (0.465) |
| *Cxcl3* | chemokine (C-X-C motif) ligand 3 | 1.44 (0.878) | 2.39 (0.269) | -1.96 (0.281) | 3.78 (0.373) | 2.04 (0.434) |
| *Il1b* | interleukin 1 beta | 1.20 (0.720) | 3.93 (0.204) | -1.62 (0.935) | 8.11 (0.262) | 3.99 (0.246) |
| *Il6* | interleukin 6 | 4.00 (0.173) | 5.64 (0.113) | 1.35 (0.513) | 13.2 (0.328) | 15.9 (0.156) |
| *Il10* | interleukin 10 | 2.26 (0.225) | 1.78 (0.342) | 3.27 (0.068) | 6.07 (0.127) | **8.38 (0.026)** |
| *Il12a* | interleukin 12a | 1.48 (0.530) | 1.89 (0.302) | 1.76 (0.318) | 1.71 (0.377) | -2.12 (0.449) |
| *Il12b* | interleukin 12b | -1.41 (0.360) | -1.36 (0.373) | 1.31 (0.540) | 1.56 (0.498) | -1.98 (0.271) |
| *Il15* | interleukin 15 | **-3.59 (0.012)** | **-7.52 (0.001)** | **-3.14 (0.039)** | **-8.20 (0.006)** | **-6.23 (0.006)** |
| *Ifng* | interferon gamma | 1.21 (0.906) | 2.27 (0.340) | 1.06 (0.991) | 1.62 (0.616) | 1.52 (0.562) |
| *Csf2* | colony stimulating factor 2 (granulocyte-macrophage) | -10.10 (0.374) | -3.32 (0.288) | 1.89 (0.428) | -4.35 (0.377) | -2.02 (0.380) |
| *Vegfa* | vascular endothelial growth factor A | -1.32 (0.397) | **-4.86 (0.032)** | 1.08 (0.901) | -1.26 (0.506) | -3.82 (0.072) |
| *Tnf* | tumor necrosis factor | -1.05 (0.658) | -1.07 (0.823) | -1.26 (0.887) | -2.69 (0.520) | -1.41 (0.315) |
| *Tgfb2* | transforming growth factor, beta 2 | 1.56 (0.059) | **2.32 (0.049)** | **1.11 (0.031)** | -1.20 (0.659) | **1.82 (0.031)** |

Data shown are expressed as the fold-regulation in transcript abundance relative to 5-day sham MK2^+/+^. Fold-regulation: fold-change values greater than one indicate an increase in transcript abundance, relative to that of sham MK2^+/+^ hearts, and the fold-regulation is equal to the fold-change. Where the transcript abundance is less than that of sham MK2^+/+^ hearts, the fold-change is less than one and the fold-regulation is the negative inverse of the fold-change. *P*-values are indicated in parentheses.
